# Cell-autonomous immune dysfunction driven by disrupted autophagy in *C9orf72*-ALS iPSC-derived microglia contributes to neurodegeneration

**DOI:** 10.1101/2022.05.12.491675

**Authors:** Poulomi Banerjee, Arpan R Mehta, Raja S Nirujogi, James Cooper, Owen G James, Jyoti Nanda, James Longden, Karen Burr, Andrea Salzinger, Evdokia Paza, Judith Newton, David Story, Suvankar Pal, Colin Smith, Dario R Alessi, Bhuvaneish T Selvaraj, Josef Priller, Siddharthan Chandran

## Abstract

The most common genetic mutation found in familial and sporadic amyotrophic lateral sclerosis (ALS), as well as fronto-temporal dementia (FTD), is a repeat expansion in the *C9orf72* gene. *C9orf72* is highly expressed in human myeloid cells, and although neuroinflammation and microglial pathology are widely found in ALS/FTD, the underlying mechanisms are poorly understood. Here, using human induced pluripotent stem cell-derived microglia-like cells (hiPSC-MG) harbouring *C9orf72* mutation (mC9-MG) together with gene-corrected isogenic controls (isoC9-MG) and C9ORF72 knock-out hiPSC-MG (C9KO-MG), we show that reduced C9ORF72 protein is associated with impaired phagocytosis and an exaggerated inflammatory response upon stimulation with lipopolysaccharide, driven by sustained activation of NLRP3 inflammasome and NF-κB signalling. Analysis of the hiPSC-MG C9ORF72 interactome revealed an association of C9ORF72 with key regulators of autophagy, a process involved in the homeostatic regulation of the innate immune response. We found impaired initiation of autophagy in C9KO-MG and mC9-MG. Furthermore, through motor neuron-microglial (MN-MG) co-culture studies, we identified that autophagy deficit in mC9-MG led to increased vulnerability of C9 MNs to excitotoxic stimulus. Pharmacological activation of autophagy ameliorated the sustained activation of NLRP3 inflammasome and NF-κB signalling, reversed the phagocytic deficit found in mC9-MG and also reduced MN death in MN-MG co-cultures. We validated these findings in blood-derived macrophages from people with *C9orf72* mutation. Our results reveal an important role for C9ORF72 in regulating microglial immune homeostasis and identify dysregulation in human myeloid cells as a contributor to neurodegeneration in ALS/FTD.

**Teaser:** Disrupted autophagy led immune activation in microglia results in enhanced motor neuronal death in *C9orf72*-ALS.

## Introduction

ALS is a devastating and incurable neurodegenerative disease characterised by selective loss of motor neurons (*1*). Although the aetiology of ALS is largely unknown, the finding that familial ALS (fALS) is clinically indistinguishable from sporadic ALS and shares a common pathology of TDP-43 proteinopathy, supports the study of monogenetic causes to better understand the unifying pathogenic mechanisms in order to develop effective treatments (*2*). For instance, hexanucleotide repeat expansions in the intronic region of *C9orf72* is the most common known cause of familial ALS and FTD (*3*). Proposed causative mechanisms underlying mutation in *C9orf72* are loss of function of C9ORF72 protein and / or gain of toxicity through RNA foci and dipeptide repeats (*4*).

Accumulating experimental and pathological evidence reveals that ALS is a multi-cellular disorder with involvement of not only motor neurons, but also other cell types, noting also that C9ORF72 is most highly expressed in myeloid cells (*5–7*). Microglia, the resident immune cell of the CNS, are implicated in ALS with evidence from autopsy *C9orf72*-ALS patient samples, suggesting chronic activation and dysregulation of the innate immune response (*8, 9*). Microglia can be activated by a diverse range of stimuli, including, for example, neuronal damage signals released following excitotoxicity-mediated motor neuronal injury, a key proposed causal pathway in ALS (*10–12*). Indeed, the finding of the immune regulatory function of C9ORF72 through suppression of STING activity opens the possibility that reduced elimination of immune responders such as STING in C9ORF72-deficient conditions might be a driver of a dysregulated immune state in ALS (*13, 14*).

Autophagy plays an important role in regulating inflammation and can be induced upon stimulation of toll-like receptors (TLRs) (*15*). Immune function is modulated through selective degradation and or/disassembly of key inflammatory mediators, such as IKKs-activator for NF-κB pathway, NLRP3-inflammasome complex and STING (*16, 17*). Furthermore, loss of autophagy-associated proteins, such as Atg7, Atg5 and beclin, in macrophages results in enhanced production of pro-inflammatory cytokines in the presence of inflammasome activators, such as lipopolysaccharide (LPS) and nigericin (*18, 19*). Indeed impaired autophagy in myeloid cells is reported to correlate with increased inflammation and exacerbation of neuronal damage in experimental models of neurodegeneration, including in Aβ injected mice and mice expressing WT human α-synuclein (*20–22*).

Against this background, using patient-derived microglial like cells from hiPSCs and macrophages, we examined the cell-autonomous and non-cell-autonomous consequences of *C9orf72* mutation.

### Results

### mC9-MG exhibit phagocytic defect and a hyperactive phenotype following LPS stimulation

To understand the cell-autonomous consequence of *C9orf72* mutation on human microglia, we first generated monocultures of hiPSC-MG, employing an established protocol, from two pairs of *C9orf72* ALS patient iPSC lines (m*C9-1 and -2*-MG) and their paired gene-edited isogenic controls (iso*C9-1 and -2*-MG) (*23*) (Fig 1A). Briefly, embryoid bodies made from iPSCs were treated with MCSF and IL-3 to generate myeloid precursors expressing CD11b, CX3CR1 and CD45 (Suppl. Fig 1A-D). These myeloid precursors were then exposed to IL-34, GM-CSF and neural precursor cell conditioned media (NPC-CM) to generate hiPSC-MGs expressing markers for human microglia, such as TMEM119, P2Y_12_ and PU.1 (Fig 1B, Suppl. Fig 1E-H). To assess the impact of *C9orf72* mutation on microglial function, we measured the phagocytic and inflammatory response to external stimulus (Fig 1C). Phagocytosis assay revealed significantly fewer internalised pH-sensitive zymosan beads, resulting from a reduction in the rate of internalisation at every time point assessed in m*C9*-MG compared to iso*C9*-MG (Fig 1D-E). We next quantified microglial production of pro-inflammatory cytokines (IL-6 and IL-1β) in response to LPS stimulation. Although the levels of IL-6 and IL-1β were comparable under basal condition across m*C9* and iso*C9*-MGs, significantly elevated levels of IL6 and ILβ were evident in m*C9-*MG upon treatment with LPS compared with iso*C9*-MG (Fig 1F, G). These findings show that *C9orf72* mutation leads to impaired microglial phagocytosis and a hyperactivated immune phenotype following LPS stimulation.

**Fig 1:**
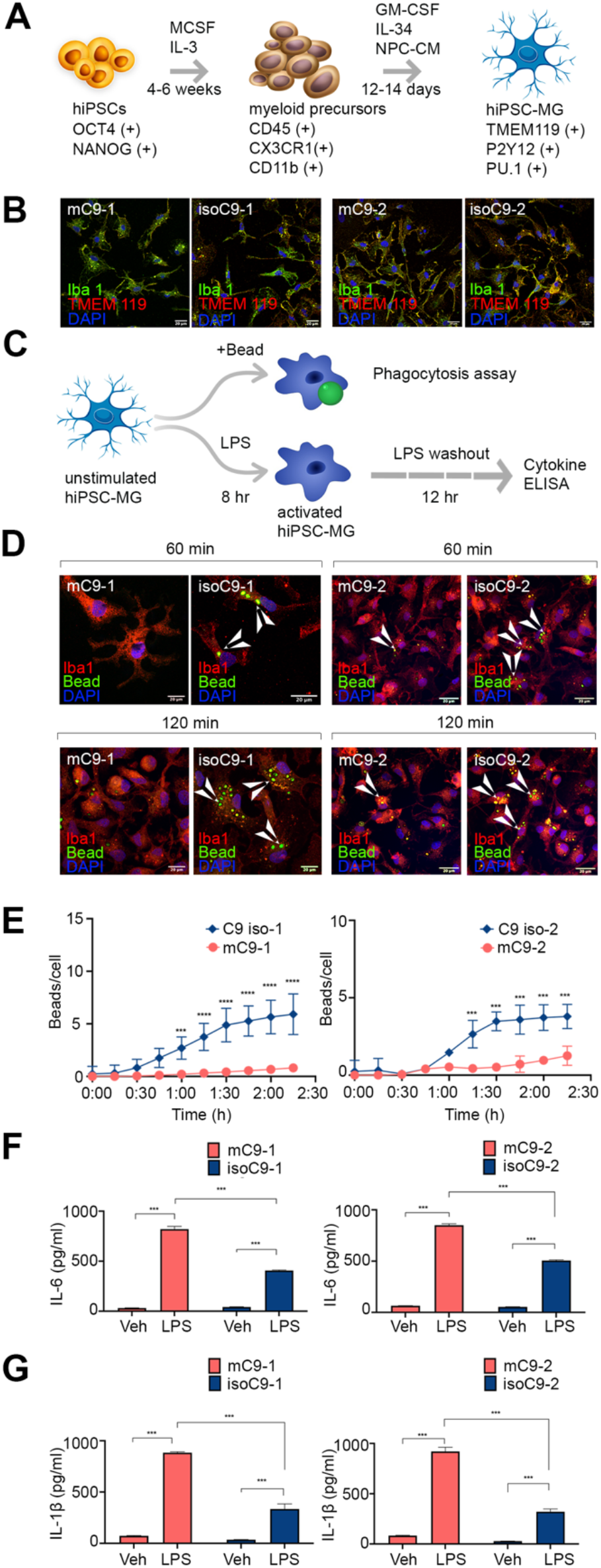
mC9-MGs display impaired phagocytosis and heightened immune response following LPS stimulation. (A) Schematic depicting stages, timeline, and key markers for the differentiation of microglia-like cells (hiPSC-MG) from human iPSCs. (B) Representative immunofluorescence images of mC9-MG and isoC9-MG demonstrating comparable staining of microglial markers: TMEM119 (red) and Iba-1 (green); scale bar = 20 μm. (C) Schematic overview of the experimental paradigm for the assessment of microglial function - phagocytosis and immune response following LPS stimulation. (D) Representative images of immunofluorescence staining showing phagocytosis assay performed with pH sensitive zymosan beads for two pairs of mC9-MG and isoC9-MG at 60 and 120 minutes (Iba-1 (red), zymosan beads (green)) scale bar = 20 μm, white arrow heads indicate phagocytosed beads (E) Graph showing real-time imaging of zymosan bead uptake at 15 minute intervals, demonstrating a phagocytic deficit in mC9-MG. Statistical analysis was performed across mC9-MG and isoC9-MG using two-way ANOVA and Tukey’s multiple comparison test, data are represented as mean +/−SEM; N=3 (*** p ≤ 0.001; **** p ≤ 0.0001). (F-G) Graph showing increased production of IL-6 (F) and IL-1β (G) in mC9-MG (red) compared to isoC9-MG (blue), suggesting an exaggerated immune response in mC9-MG following LPS stimulation; cells were treated with either LPS or 1X PBS (Vehicle control (veh)), Statistical analysis was performed using 2-way ANOVA and Tukey’s multiple comparison test, data are represented as mean +/−SD; N=3 (*** p ≤ 0.001)

### C9ORF72 interactome reveals association with regulators of autophagy in human iPSC derived microglia

C9ORF72 protein is highly expressed in myeloid cells (*5–7*), a finding that we also confirmed by quantitative immunoblot of control human iPSC-derived microglia, neurons and astrocytes (Suppl. Fig 2 A, B). Immunoblot of m*C9*-MG compared to iso*C9*-MG revealed significantly reduced abundance of C9ORF72 protein, a finding comparable to previous reports (*24*) (Fig 2A). To further investigate the impact of reduced C9ORF72 protein on microglial function, we generated microglia from a *C9orf72* null iPSC line. The *C9orf72* null iPSC line was generated from a control iPSC line using CRISPR/Cas9 genome editing technology (Suppl. Fig 3A,B,). Immunoblot of C9KO-MG confirmed the absence of C9ORF72 protein (Fig 2B,2C) compared to the parental control iPSC-derived microglia (ctrl-MG). C9KO-MG also exhibited reduced phagocytosis (Fig 2D, 2E) and increased production of IL-1β following LPS stimulation (Fig 2F), thereby phenocopying the microglial functional deficits found in m*C9*-MG. These findings suggest that reduced abundance of C9ORF72 protein contributes to the functional deficits seen in m*C9*-MG. To better understand the function and the pathways that are disrupted due to loss of C9ORF72 in hiPSC-MGs, we next evaluated the interactome of C9ORF72 in ctrl-MG. A C9ORF72 reporter iPSC line was generated by introducing an EGFP tag at the N-terminus of the C9ORF72 protein (preserving its endogenous promoter) in order to perform GFP-Trap immunoprecipitation (Suppl. Fig 4A). The EGFP-C9 line was differentiated into C9GFP-MG (Suppl. Fig 4C) and immunoblot confirmed the presence of EGFP-C9ORF72 fusion protein (Suppl. Fig 4B). Following immunoprecipitation using GFP-trap beads, the cell lysate was subjected to an unsupervised proteomic screen to identify the interactive partners of EGFP-C9ORF72 by tandem mass spectrometry. Applying stringent pre-test criteria of >2.0-fold enrichment and 1% false discovery rate (FDR), 254 potential interactors proteins, including novel partners such as FMR1, MMP9, UFL1 were identified (Fig 2G, Supplementary table 2). STRING network analysis was next performed on the top 50 proteins (GFP vs control fold enrichment >= 3.0). The 5 most significant gene ontology (GO) processes (p>0.05) identified were (i) regulation of TORC1 signalling (ii) regulation of autophagosome assembly (iii) vesicle mediated transport (iv) regulation of cell migration and (v) cellular response to stress (Fig 2H). Amongst these, autophagy-associated proteins, such as SMCR8 and WDR41, were enriched more than 5-fold; these were further validated using western blotting (Suppl. Fig 4D). Finally, enrichment for multiple autophagy-associated RAB proteins (*25*) (RAB5, RAB8A, RAB39Ab, RAB13, RAB 34, RAB21, RAB 10) was observed in the C9ORF72 interactome. Taken together, these findings suggest an important role of C9ORF72 in the regulation of autophagy in human microglia.

**Fig 2:**
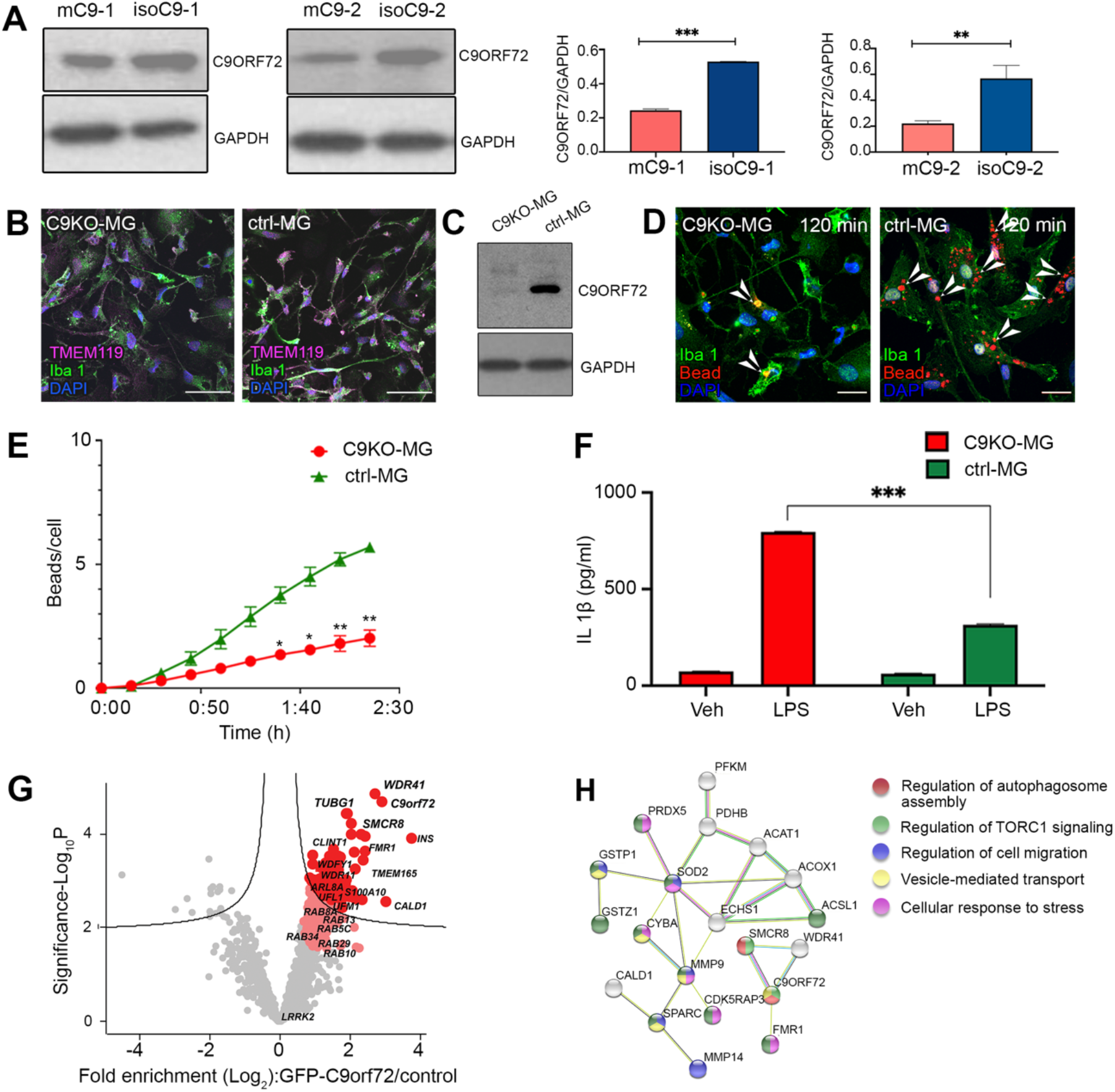
mC9-MG and C9KO-MG exhibit reduced abundance of C9ORF72 protein and mass spectrometric analysis of C9ORF72 interactome revealed its association with regulators of autophagy. (A) Immunoblot with respective densitometric analysis showing reduced abundance of C9ORF72 protein (~55kDa) across two lines of mC9-MG (red) when compared to their respective isogenics (blue), data are represented as mean +/−SD N=3. Statistical analysis was performed employing students t-test, (** p ≤ 0.01,*** p ≤ 0.001) (B) Representative images of immunofluorescence staining of TMEM119 (magenta) and Iba-1 (green) of C9KO-MG and ctrl-MG, scale bar=50 μm. (C) Immunoblot validating loss of C9ORF72 protein (55kDa) in C9KO-MG. (D) Representative images of immunofluorescence staining showing phagocytosis assay performed with pH sensitive zymosan beads for C9KO-MG and ctrl-MG at 120 minutes (Iba-1 (green), zymosan beads (red) scale bar = 20 μm; white arrow heads indicate phagocytosed beads (E) Graph showing real-time imaging of zymosan bead uptake at 15 minute intervals demonstrating a phagocytic deficit in C9KO-MG when compared to control (ctrl-MG), statistical analysis was performed using two-way ANOVA and Tukey’s multiple comparison test, data are represented as mean +/−SEM (* p ≤ 0.05; ** p ≤ 0.01), N=3; (F) Graph demonstrating increased production of 1L-1β for C9KO-MGs following LPS induction; cells were treated with either LPS or 1XPBS (Vehicle control (veh)); statistical analysis was performed using two-way ANOVA and Tukey’s multiple comparison test, data are represented as mean+/−SD; N=3 (*** p ≤ 0.001) (G) Volcano plot demonstrating enrichment of C9ORF72 interactors at 1% false discovery rate (FDR) (dark red) and 5% FDR (light red). There is significant enrichment in autophagy-associated proteins, SMCR8 and WDR41, in the C9ORF72 interactome; n=4 (H) STRING network analysis of the C9ORF72 interactors (1% FDR); nodes represent the functional enrichment of the network.

### C9orf72 loss-of-function disrupts autophagy initiation in mC9-MG

Recognising that C9ORF72 interactors SMCR8, WDR41 and Rab5 are integral to the autophagy initiation machinery (*26, 27*), we first investigated basal autophagy in mC9-MG. We measured the formation of autophagosomes in the presence of bafilomycin (preventing the fusion of autophagosomes and lysosomes) by quantitative immunostaining with the autophagosome membrane proteins LC3 and p62. Upon bafilomycin treatment, we observed significantly fewer LC3 and p62 puncta in the mC9-MG when compared to isoC9-MG (Fig 3A, B). Interestingly, C9KO-MGs, compared to ctrl-MG, also displayed significantly reduced number of p62 puncta post bafilomycin treatment (Fig 3B). Furthermore, the autophagy deficit was also confirmed by immunoblot which demonstrated reduced levels of p62 and LC3 at 2, 4 and 6 hours in presence of bafilomycin in mC9-MGs and C9KO-MGs compared to their isogenic controls and ctrl-MGs respectively (Fig 3C, Suppl. Fig 5). To explore whether this autophagic deficit reflected a deficit in the induction of autophagy, we next measured autophagy flux using lentiviral overexpression of a tandem dual fluorescent reporter (mCherry-GFP-p62)(*28*). This pH-sensitive tool enables the assessment of autophagosomes which are GFP^+ve^, mCherry^+ve^ (yellow puncta), versus autolysosomes which are GFP^-ve^, mCherry^+ve^ (red puncta). We observed significantly fewer autophagosomes in the mC9-MG compared to isogenic controls (Fig 3D). Furthermore, we found that the ratio of autolysosomes to autophagosomes was not altered in mC9-MG (Fig 3D), a finding consistent with the autophagic deficit in mC9-MG being driven by an impairment in the induction of autophagy. Interestingly, this deficit in autophagy also became evident during phagocytosis where reduced numbers of LC3 puncta were observed during the internalisation of zymosan beads in mC9-MG compared to their isoC9-MG (Suppl.Fig 6). Thus, we show deficits in autophagy in mC9-MG and C9KO-MG.

**Fig 3:**
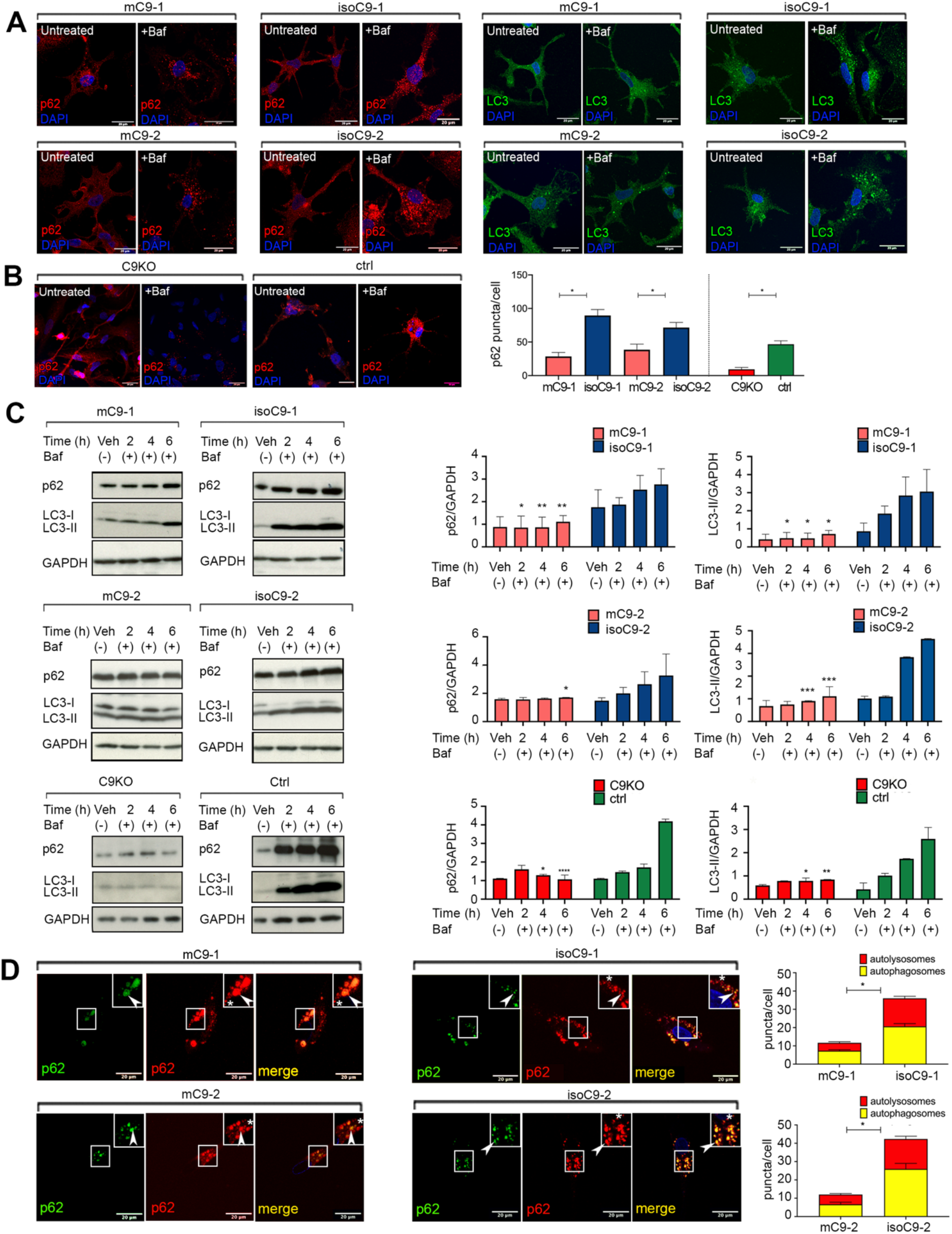
mC9-MG demonstrate a deficit in the initiation of autophagy. (A) Representative images of immunofluorescence staining of p62 (left panel) and LC3 (right panel) positive puncta after 6h of DMSO (vehicle control) and bafilomycin treatment, demonstrating fewer p62 and LC3 puncta in mC9-MG as opposed to isoC9-MG in the bafilomycin treated cells, scale bar = 20 μm (B) Representative images of immunofluorescence staining showing the reduced number of p62+ve puncta at 6h post bafilomycin treatment in C9KO-MG when compared to ctrl-MG, bar graph (right) representing the quantification for Fig 3A and Fig 3B depicting an increase in the number of p62 puncta after 6 hours of bafilomycin treatment in isoC9-MGs and ctrl-MG when compared to mC9-MGs and C9KO-MG; data is represented as mean +/−SD. Statistical analysis was performed using two-way ANOVA and Tukey’s multiple comparison test, data are represented as N=3(n=20 cells/genotype), * p ≤ 0.05 (C) Representative immunoblots (left) showing reduced turnover of p62 (~60kDa) and LC3B(ii) (~15kDa) in two pairs of mC9-MG and C9KO-MG compared to their respective isoC9-MG and ctrl-MG in the presence of bafilomycin at 2h,4h,6h demonstrating an impairment in autophagy initiation. Bar graphs (right) represent the densitometric quantification of p62 and LC3B(ii) normalised to loading control GAPDH; data are represented as mean+/−SD N=3. Statistical analysis was performed across mC9-MG, isoC9-MG and C9KO-MG, ctrl-MG at 2h,4h, 6h time point using two-way ANOVA and Sidak’s multiple comparison test, * p < 0.05, ** p < 0.01, *** p < 0.001, **** p ≤ 0.0001 (D) Representative images from live imaging of mcherry-EGFP-p62 dual reporter probe transduced in 2 pairs of mC9-MG and isoC9-MG; the white arrowhead in the inset represents autophagosomes (GFP^+ve^, mCherry^+ve^) and the white asterisk represents autolysosomes (GFP^-ve^mCherry^+ve^). Stacked bar graphs (right) demonstrating the quantification of the autophagosomes and autolysosomes. These show: (i) a reduction of number of autophagosomes (ii) the ratio of the autophagosomes to autolysosomes of mC9-MG and their respective isoC9-MG. Statistical analysis was performed using two-way ANOVA and Tukey’s multiple comparison test for the number of autophagosomes across mC9-MG and isoC9-MG, data are represented as mean+/−SD; N=3 {n(mC9-1=20cells, mC9-2=18, isoC9-1=20, isoC9-2=20)}, * p ≤ 0.05.

### Disrupted autophagy leads to sustained activation of NLRP3-inflammasome and NF-κB signalling in mC9-MG

Since autophagy regulates the attenuation of pro-inflammatory signalling (*29*), we next investigated the impact of *C9orf72* mutation on the NLRP3-inflammasome and NF-κB pathway in m*C9*-MGs. We treated m*C9*-MG and iso*C9*-MG in a time-limited exposure (8h) with LPS in order to stimulate an immune response. Following this, we removed LPS and monitored the post-washout time-course through the collection of cellular lysates at 4h, 8h, 12h (Fig 4A). Following the removal of LPS stimulation, we observed a comparable increase in the levels of NLRP3 in m*C9*-MG, iso*C9*-MG and ctrl-MG at the first four hours post LPS washout, indicating a comparable inflammatory response to LPS stimulation (Fig 4B, Suppl. Fig 7 A, B). However, in ctrl-MG and iso*C9*-MG, levels of NLRP3 returned to near-basal levels between 8-12 hours (Fig 4B, Suppl. Fig 7 A, B). This reduction in NLRP3 protein was accompanied by a reciprocal increase in p62 levels, marking the activation of autophagy, over the same time interval (Fig 4B, Suppl. Fig 7 A, C) in ctrl-MG and iso*C9*-MG. In contrast, in m*C9*-MG, there was no reduction in NLRP3, and significantly reduced levels of p62 (Fig 4B) at 4, 8 and 12 h following LPS removal. These findings are consistent with mutation-dependent impairment in the autophagy initiation machinery, resulting in the sustained activation of NLRP3-inflammasome following LPS induction.

**Fig 4:**
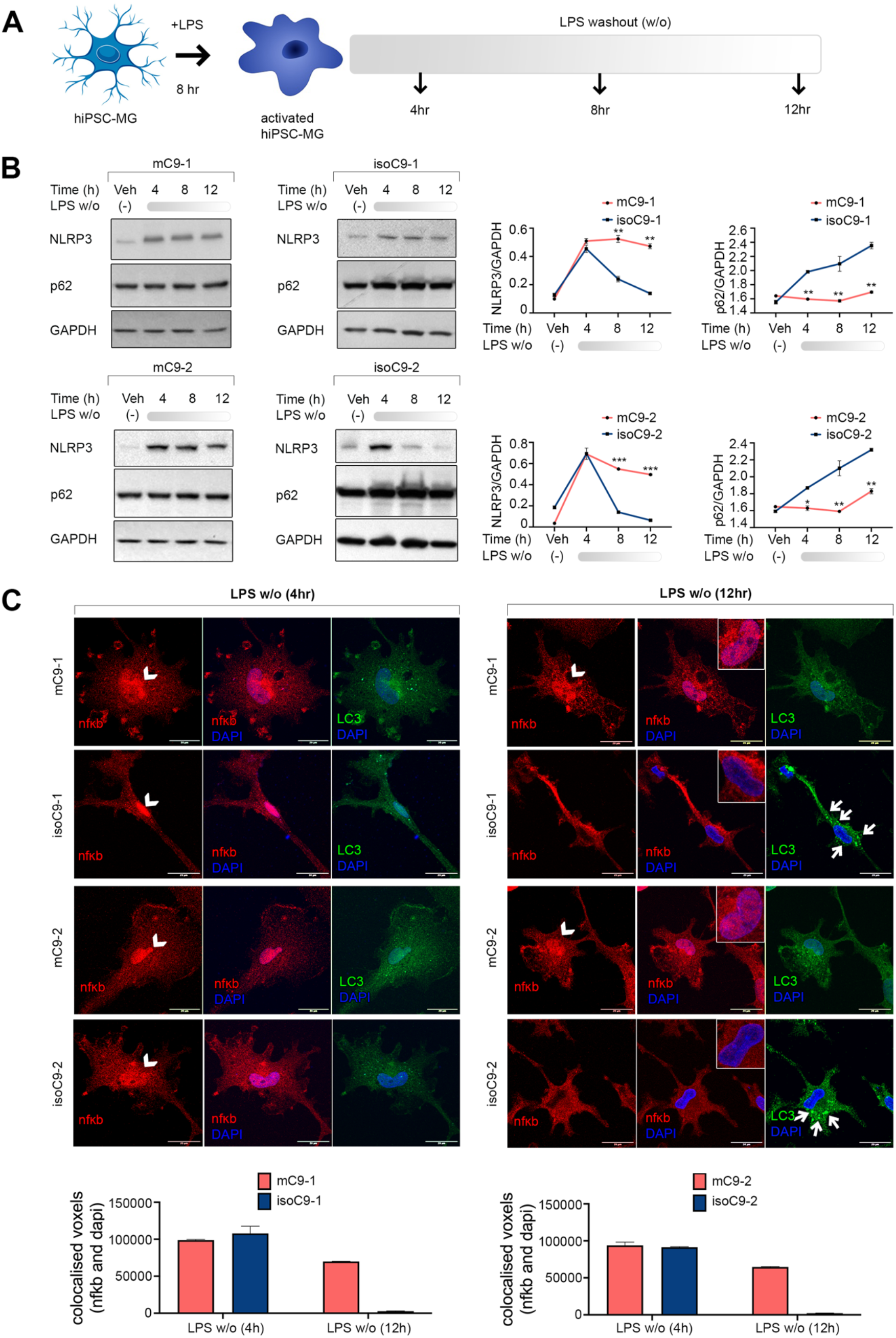
Disrupted autophagy in mC9-MG results in inadequate attenuation of inflammasome activation and NF-κB signalling following LPS stimulation. (A) Schematic of the experimental paradigm for the assessment of the dynamics of NLRP3 inflammasome and NF-κB signalling following LPS stimulation. mC9-MG and isoC9-MG were treated with 100 ng/ml of LPS for 8h and, following LPS removal, whole cell lysates were harvested at 4h, 8h and 12h; cells were treated with either LPS or 1XPBS (Vehicle control (veh)) (B) Immunoblots showing differential immunoreactivity for NLRP3 (~110kDa), p62 (~60kDa) at 4h, 8h and 12h post LPS washout, densitometric values for NLRP3 and p62 were normalised to GAPDH and line graphs for isoC9-MGs (blue) show a gradual decline of NLRP3 with a concomitant reciprocal increase in p62 indicative of autophagy induction. Conversely the mutant cells (red line) did not show a decline in NLRP3 levels over time and show no evidence of a time dependent increase in p62 levels. Data are represented as mean +/−SD for 3 independent experiments, statistical analysis across mC9-MG and isoC9-MG at 4h, 8h and 12h was performed using two-way ANOVA and Sidak’s multiple comparison test * p < 0.05, ** p < 0.01, *** p < 0.001. (C) Representative images of immunofluorescence staining for NF-κB and LC3 demonstrating NF-κB signalling and autophagy activation respectively at 4h and 12h post LPS washout. At 4h (left panel) both mC9-MG and isoC9-MG show nuclear localisation of NF-κB indicative of activated NF-κB signalling. At 12h, mC9-MG demonstrate sustained nuclear localisation and activation of NF-κB signalling (right panel and the insets) as opposed to the disappearance of NF-κB from the nucleus in the isoC9-MG suggesting attenuation of NF-κB signalling. Note the appearance of LC3 puncta in the isoC9-MG, indicative of autophagy induction as opposed to mC9-MG. The nuclear region is highlighted using white arrow heads and the presence of LC3 puncta are indicated using small white arrows at the 12 h time point. Graphs at the bottom show the quantification of co-localised voxels of NF-κB and DAPI which is maximal at 4 hours and declines over 12 hours in the isogenic control-a trend that was not found in mC9-MG, data is represented as mean +/−SD, data represents n = 20 cells across all genotypes.

We next investigated NF-κB pathway dynamics, another inflammatory pathway reliant on autophagy for negative regulation (*30*). Four hours after LPS removal, in both m*C9* and their isogenic pairs, we observed nuclear localisation of NF-κB (Fig 4C), a finding that implies inflammation-induced activation of this pathway, noting that, under basal conditions NF-κB is largely localised to the cytoplasm (Suppl. Fig 8). This activated state as evidenced by nuclear localisation exhibited a tendency to return to basal state (cytoplasmic localisation) in the iso*C9*-MG and ctrl-MG at 12 hours (Fig 4C, Suppl. Fig 7D, E). In contrast, m*C9*-MG displayed sustained nuclear localisation of NF-κB at 12 hours post-LPS washout (Fig 4C). Collectively, these data suggest that *C9orf72* mutation leads to the sustained activation of both NLRP3 and NF-κB signalling in iPSC-MG following LPS stimulation.

### Pharmacological activation of autophagy ameliorates the impaired phagocytosis and hyperactive state of mC9-MG and C9KO-MG

To determine whether boosting autophagy could ameliorate both the phagocytic deficit and activated immune state in m*C9*-MGs, we next treated m*C9*-MGs with rapamycin, a pharmacological activator of autophagy (*31*) (Fig 5A). Rapamycin treatment resulted in a significant increase in p62-positive puncta in bafilomycin-treated m*C9*-MG (Suppl. Fig 9A,B) and C9 KO-MG (Suppl. Fig 9C,D). This finding was further validated by western blot analysis, demonstrating a time dependent increase in p62 level for m*C9*-MGs, iso*C9*-MGs and C9KO-MG, ctrl-MG and LC3(II) levels for mC9-MG and isoC9-MG following rapamycin treatment in presence of bafilomycin, reflective of autophagic induction (Suppl. Fig 9 E-J). We next challenged m*C9*-MGs, iso*C9*-MGs and ctrl-MGs with LPS in the presence of rapamycin to investigate the impact on NLRP3 and p62. Notably, we observed a significant reduction in NLRP3 immunoreactivity at a steady-state following LPS stimulation in the m*C9*-MG, iso*C9*-MG and ctrl-MG, with a reciprocal significant increase in p62 level compared to m*C9*-MGs that were not treated with rapamycin at 12h (Fig 5B, Suppl. Fig10B). This rapamycin-induced amelioration of the hyper-activated state of m*C9*-MG was also found in the NF-κB signalling pathway, which demonstrated a reduction in nuclear localisation of NF-κB and an increase in the appearance of LC3 puncta, indicative of autophagy-mediated suppression of NF-κB signalling (Fig 5C, D, Suppl. Fig 10A, 10D). Next, we determined whether boosting autophagy in m*C9*-MG and C9KO-MG reverses the functional deficits of phagocytosis and increases cytokine production. Rapamycin treatment resulted in a significant reduction in the production of the pro-inflammatory cytokines IL-6 and IL-1β in response to LPS in m*C9*-MG, C9KO-MG, isoC9-MG and ctrl-MG (Fig 5E-I Suppl. Fig 10C), and also ameliorated the phagocytic deficit in m*C9*-MG and C9KO-MGs (Fig 5J-L).

**Figure 5:**
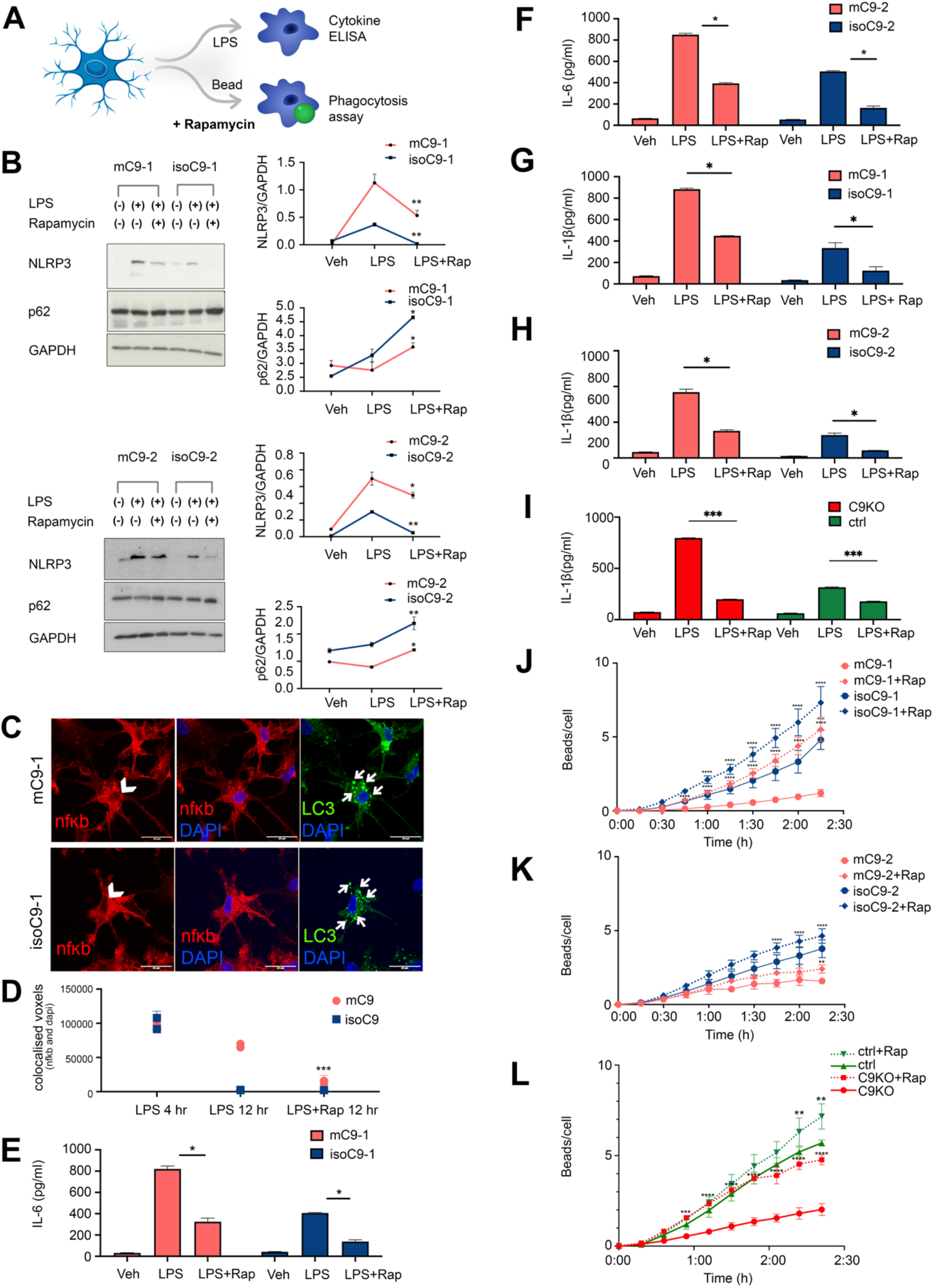
Pharmacological activation of autophagy with rapamycin ameliorates the sustained immune activation in mC9-MG and C9KO-MG. (A) Schematic showing the experimental setup wherein the cells are treated with rapamycin for 12 h, post LPS washout and during zymosan bead uptake assay, (B) Immunoblots and their quantification showing an increase in p62 (~60kDa) immunoreactivity and a reciprocal decline in NLRP3 (~110kDa) immunoreactivity across mC9-MG and isoC9-MG in presence of rapamycin. The cells were either treated with vehicle (Veh), LPS or LPS+Rapamycin (LPS+Rap). Statistical analysis was performed for mC9-MG and isoC9-MG across rapamycin treated and untreated condition in presence of LPS using two-way ANOVA and Sidak’s multiple comparison test * p < 0.05, ** p < 0.01. Data represent mean+/−SD across 3 independent experiments. (C) Representative images of immunofluorescence staining showing the reduction of nuclear localisation of NF-κB in rapamycin-treated mC9-MG, demonstrating an attenuation of NF-κB signalling. The nuclear regions are indicated using white arrow heads and concurrent increase in appearance of LC3 puncta are highlighted using white arrows (D) Shows the quantification of the co-localised voxels of NF-κB and DAPI post rapamycin treatment. data is represented as mean +/−SD; n = 20 cells across all genotypes. Statistical analysis was performed for mC9-MG and isoC9-MG across rapamycin treated and untreated condition in presence of LPS at 12 h time point using two-way ANOVA and Sidak’s multiple comparison test *** p < 0.001(E-H) Cytokine ELISA of IL-6 and IL-1β demonstrates suppression of the production of pro-inflammatory cytokines in mC9-MG following rapamycin treatment. Cells were treated with either vehicle (Veh), LPS or LPS+Rapamycin (LPS+Rap); data are represented as mean +/−SD; statistical analysis was performed using two-way ANOVA and Sidak’s multiple comparison test (* p ≤ 0.05); N=3 (I) IL-1β ELISA demonstrates the amelioration of immune response in C9KO-MGs mirroring mC9-MGs as a result of rapamycin treatment, cells were treated with either vehicle (Veh), LPS or LPS+Rapamycin, data are represented as mean +/−SD; statistical analysis was performed using two-way ANOVA and Sidak’s multiple comparison test *** p ≤ 0.001); N=3 (J,K,L) Graph showing real-time imaging of zymosan bead uptake assay demonstrating rapamycin mediated amelioration of the phagocytic deficit in mC9-1-MGs and mC9-2-MGs and C9KO-MGs, statistical analysis was performed across rapamycin treated and untreated condition for all genotypes using 2 way ANOVA and Tukey’s multiple comparison test (* p ≤ 0.05; ** p ≤ 0.01; *** p ≤ 0.001; **** p ≤ 0.0001) and error bars represents +/−SEM; N=3.

### Disrupted autophagy in mC9-MG contributes to enhanced motor neuronal death following excitotoxic insult

To next examine the functional impact of disrupted microglial autophagy on C9 mutant motor neurons (mC9-MNs), we undertook co-culture studies of C9 mutant MN and C9 mutant MG (mC9 MN-MG), isogenic C9 MN and isogenic MG (isoC9 MN-MG) and control MN and control MG (ctrl MN-MG) (Fig 6A,B, Suppl. fig 11A). Spinal MNs were generated using an established protocol (*11, 32*). We observed comparable survival of MNs across mC9 MN-MG, isoC9 MN-MG and ctrl MN-MG under basal conditions (Fig 6 B, Suppl. fig11A). We next treated these co-cultures with AMPA to model excitotoxicity that we and others have previously shown to be selectively toxic to isolated mC9 MNs (*11, 24, 33*). Noting also that neuronal excitotoxicity has been shown to induce microglial activation, we predicted that mC9 MN-MG co-culture will show increased MN death compared to isolated MN cultures alone following 24h treatment with 100μM AMPA (*12*). First, we examined the effect of AMPA on microglial activation state on isolated MG and separately in co-culture with MNs. Using IL-1β as a marker of activated microglia we showed that AMPA does not elicit microglial activation in isolated MG cultures (mC9, isoC9, ctrl) (Suppl. Fig 11B). However, we observed significantly increased activation of mC9-MG in the mC9MN-MG co-cultures when compared to isoC9 MG in isoC9MN-MG and ctrl MG in ctrl MN-MG co-cultures as reflected by IL-1β level (Fig.6C, Suppl. Fig 11C). Next, we assessed survival of MNs (mC9, isoC9, ctrl) in isolation and separately when co-cultured with respective genotype MGs upon AMPA stimulation. As previously reported, isolated mC9MN cultures revealed significantly increased MN death compared to isoC9MN, (mC9-41±4.72 % vs iso C9-29±1.73%) and ctrl MNs following AMPA stimulation (Fig 6D, Suppl. Fig 11D). AMPA treatment of mC9 MN-MG compared to isoC9 MN-MG and ctrl MN-MG co-cultures revealed a significantly enhanced mC9 MN death compared to isoC9-MNs (mC9 61% ±3.46%vs isoC9 34%±1%) and ctrl MNs, suggestive of additive neurotoxic impact of mC9-MG on mC9-MN. (Fig 6E, Suppl.Fig 11E). In contrast MG numbers were comparable between the mutant versus iso co-cultures and ctrl co-cultures (Fig 6F, Suppl. Fig 11F). Together, these findings suggest the increased activation of mC9MG following AMPA exposure in co-cultures is neuronally driven and leads to enhanced mC9MN death.

**Figure 6:**
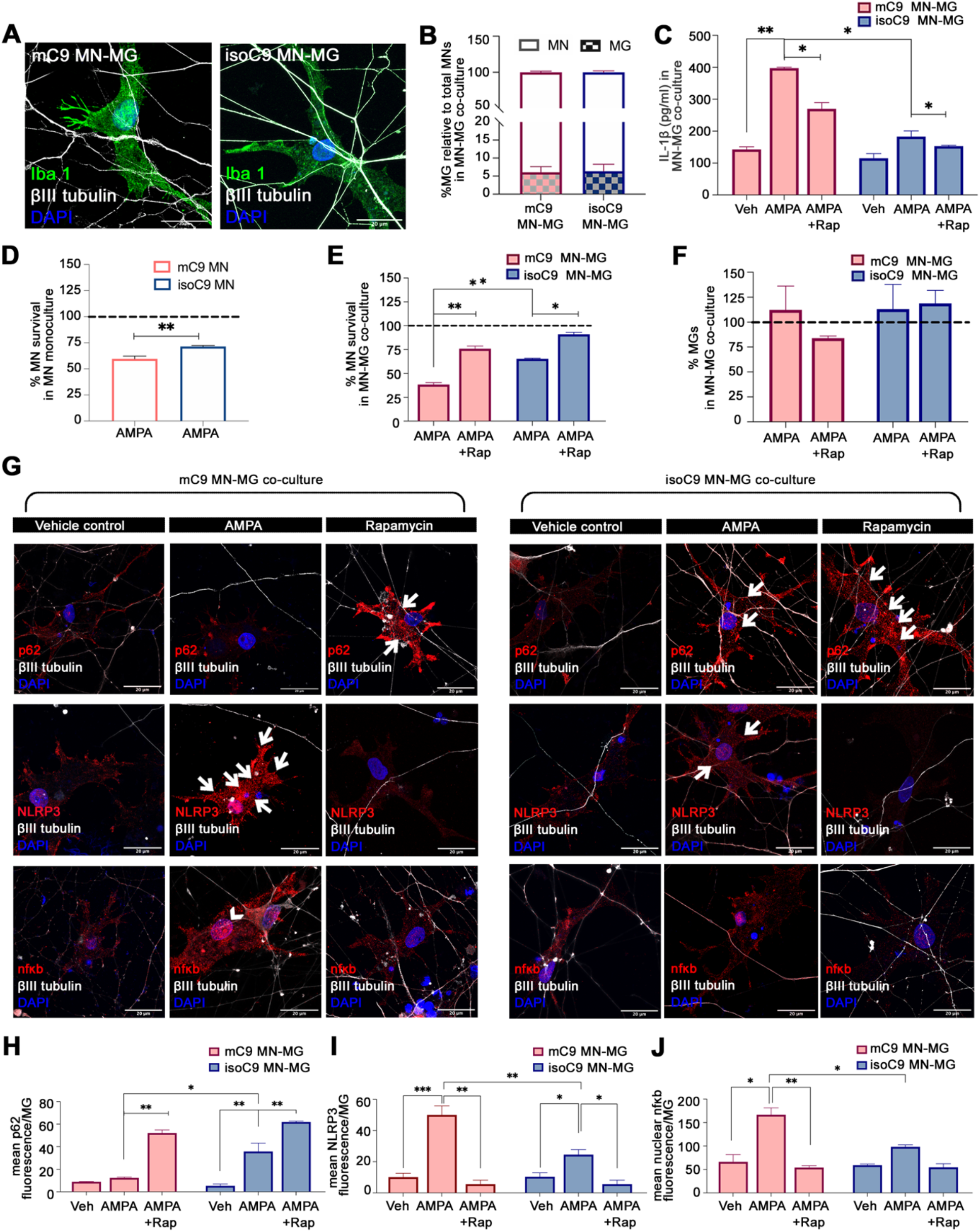
Disrupted autophagy in mC9-MG contributes to enhanced motor neuronal death following excitotoxic insult. (A) Representative immuno-fluorescence images for Iba1 (green) and βIII tubulin (grey) in mC9 motor neuron-microglial (mC9 MN-MG) co-culture and isoC9 motor neuron-microglial (isoC9 MN-MG) co-culture. Scale bar=20μm (B) Graph representing percentage of microglia (MG) relative to total motor neurons (MN) in mC9 MN-MG co-culture and isoC9 MN-MG co-culture. Data are represented as mean +/−SD N=3 (C) Graph representing the production of IL1β in mC9 MN-MG and isoC9 MN-MG co-cultures in vehicle treated condition and following AMPA challenge in absence and presence of rapamycin. Data are represented as mean +/−SD N=3 (D) Graph representing the percentage of motor neuronal survival in mC9-MN monoculture and isoC9-MN monoculture relative to their vehicle treated controls (represented by dashed line) following AMPA treatment for 24 hours. Data are represented as mean +/−SD N=3 (E) Graphs showing the percentage survival of motor neurons in mC9 MN-MG co-culture and isoC9 MN-MG coculture following AMPA treatment for 24 hours in absence and presence of rapamycin relative to their vehicle treated controls as represented by the dashed line. Data are represented as mean +/−SD N=3 (F) Graph representing the percentage survival of MGs in mC9 MN-MG co-culture and isoC9 MN-MG co-culture relative to their vehicle treated condition (represented by the dashed line) following AMPA treatment for 24 hours in absence and presence of rapamycin. Data are represented as mean +/−SD N=3 (G) Representative images of fluorescence staining for microglial p62/NLRP3/nfκb across vehicle treated, AMPA treated conditions in absence and presence of rapamycin in mC9 MN-MG co-culture and isoC9 MN-MG co-culture. White arrows indicate the cytoplasmic localisation of p62 and NLRP3 staining and arrow heads indicate the nuclear localisation of nfkb staining Scale bar=20μm (H,I,J) Graph representing the quantification of the mean fluorescence intensity of p62/NLRP3/nfκb per microglial cell across vehicle treated, AMPA treated conditions in absence and presence of rapamycin in mC9 MN-MG and isoC9 MN-MG co-cultures. 20 microglial cells from three biological replicates have been analysed per condition against each genotype, data are represented as mean +/−SD N=3.

To next test whether microglia mediated increased neuronal injury following AMPA treatment reflects sustained immune activation due to disrupted autophagy we next measured NLRP3, NF-κB and autophagy marker p62 in co-cultured microglial cells following AMPA stimulus. Increased NLRP3 and nuclear NF-κB alongside a concomitant reduction of p62 level was seen in mC9-MG compared to isoC9-MGs and ctrl-MGs in the MN-MG co-cultures following AMPA treatment (Fig 6G, Suppl. Fig 11 G-I). To test whether boosting autophagy would reduce MN death, we treated isolated MN and co-cultures of MN-MG with rapamycin. Rapamycin treatment resulted in reduced IL-1β and mC9 MN death in mC9-MN/MG co-culture, but not of isolated MN cultures (Fig. 6C, 6E, Suppl. Fig 11C, E, J). Together, these findings show that disrupted microglial autophagy in mC9-MG contributes to MN death following neuronal excitotoxicity and that boosting microglial autophagy reduced motor neuron death.

### Blood-derived macrophages from *C9orf72* carriers display an activated phenotype and impairment of phagocytosis

To further assess the disease relevance of our iPSC findings, we next directly studied blood derived macrophages from volunteers with the *C9orf72* mutation, and their respective age and sex-matched controls (Table 1). We isolated peripheral blood mononuclear cells and differentiated them into macrophages by treating them with 80ng/ml MCSF for 7 days (CD45+ve and Iba1+ve) (Fig 7A). Quantitative immunoblot revealed reduced abundance of C9ORF72 in C9 mutation-derived macrophages (Fig 7B, C). Concomitantly, we also observed significantly higher levels of NLRP3 protein in C9 macrophages (Fig 7B, C), when compared to their respective age- and sex-matched controls. C9 macrophages also secreted significantly elevated levels of pro-inflammatory cytokines, IL-6 and IL-1β, following LPS treatment (Fig 7G-H). We next examined phagocytosis; macrophages from C9 positive subjects demonstrated a phagocytic deficit with significantly reduced zymosan bead uptake over a period of 2 h (Fig 7D,F). Significantly, induction of autophagy in the C9 macrophages with rapamycin resulted in reduced IL-6 and IL-1β as well as amelioration of the phagocytosis phenotype (Fig 7 F,G-H). Collectively these findings are consistent with a mutation-dependent dysregulation of autophagy in macrophages and MG that results in an altered immune activation status that is reversible.

**Table 1:** Clinical demographics of cases and controls for the PBMC-derived macrophage studies. F = female, M = male; UL = upper limb, LL= lower limb.

| ID | Genotype | Diagnosis | Sex | Ethnicity | Age of onset (years) | Age at blood sample (years) | Disease duration (months) | Site of onset | Drug history | Family history |
| --- | --- | --- | --- | --- | --- | --- | --- | --- | --- | --- |
| Con-1 | Control | - | F | Caucasian | - | 51 | - | - | Nil | Nil |
| Con-2 | Control | - | M | Caucasian | - | 42 | - | - | Nil | Nil |
| Con-3 | Control | - | F | Caucasian | - | 78 | - |  | Thyroxine | Nil |
| C9 ALS-1 | C9orf72 | ALS | F | Caucasian | 52 | 54 | 24 | LL | Riluzole | FTD |
| C9 ALS-2 | C9orf72 | Asymptomatic Carrier | M | Caucasian | - | 44 | - | - | Nil | ALS/FTD |
| C9 ALS-3 | C9orf72 | FTD | F | Caucasian | 76 | 77 | 84 | - | Amlodipine, atorvastatin | FTD |

**Figure 7:**
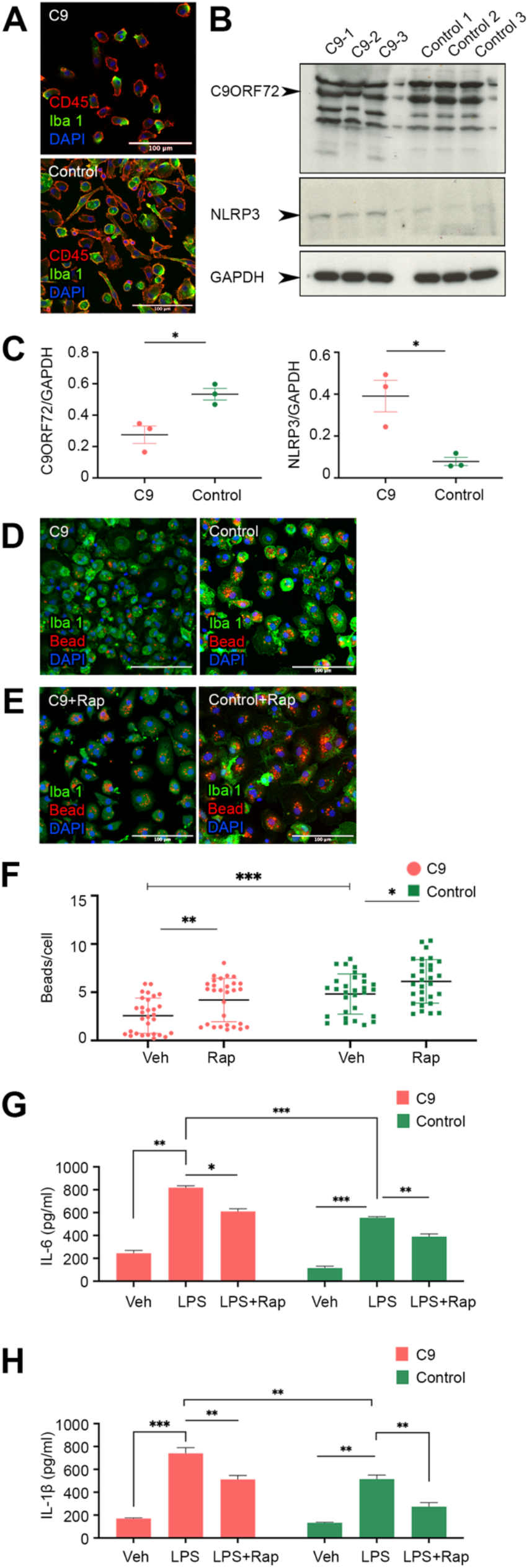
Blood-derived macrophages isolated from people with C9orf72 mutation display a hyper-active state and phagocytic deficit, recapitulating deficits observed in mC9-MGs. (A) Representative images of immunofluorescence staining demonstrating comparable staining of macrophage markers (CD45 (red) and Iba-1 (green)) in the patient and control PBMC derived macrophages. (B) Immunoblot and (C) densitometric quantification demonstrating reduced abundance of C9ORF72 (~55kDa) (graph-left), and increased levels of NLRP3 (~110kDa) (graph-right), in three patient-derived macrophage samples compared to their age- and sex-matched controls, data are represented as mean +/−SD and statistical analysis was performed using student’s t test, * p ≤ 0.05 (D) Representative images of immunofluorescence staining show phagocytic deficit in C9 ALS patient blood derived macrophages when compared to their age- and sex-matched controls (E) Representative immunostaining and quantification (F) of the number of zymosan beads post 2 hours demonstrate a significant increase in the phagocytic ability of rapamycin treated C9 ALS patient derived blood macrophages, statistical analysis was performed using 2 way ANOVA and Tukey’s multiple comparison test (* p ≤ 0.05; ** p ≤ 0.01; *** p ≤ 0.001) and data represent mean +/−SD across the pooled data from three C9-ALS cases with age/sex matched control (G) ELISA for pro-inflammatory cytokines demonstrates increased production of IL6 and IL-1β in *C9*ALS-patient derived macrophages compared to age- and sex-matched controls in unstimulated and LPS stimulated condition, thus mirroring hyper responsive behaviour of mC9-MGs following LPS challenge. Rapamycin treatment was shown to suppress the hyper responsive nature of C9 ALS macrophages in presence of LPS stimulation, statistical analysis was performed using two-way ANOVA and Tukey’s multiple comparison test (* p ≤ 0.05; ** p ≤ 0.01; *** p ≤ 0.001; **** p ≤ 0.0001) and data are represented as mean +/−SD.

## Discussion

C9ORF72 is expressed in human myeloid cells, yet it is perhaps surprising that its function in human immune cells has only been partially explored (*13, 34*). Through the combination of patient iPSC-derived microglia, isogenic controls and patient-derived macrophages, we show that mutant myeloid cells are deficient in phagocytosis and are hyper-reponsive to inflammatory stimuli. Further, employing a co-culture of motor neuron and microglia, we also demonstrate that C9ORF72 mutation confers a cell-autonomous deficit in autophagy leading to a hyper-activated NF-κB and NLRP3 axis, which plays a contributory role in eliciting enhanced motor neuronal death, thereby providing new insights into the mechanism and consequence of dysregulated immune response found in *C9orf72*-ALS.

Multiple lines of evidence including p62^+^/ TDP-43 inclusions in patient autopsy samples and discovery of a number of ALS causal genes implicated in the autophagy pathway suggest an important role for autophagy dysregulation in ALS (*35*). C9ORF72 has also been implicated in autophagy, although the precise mechanism of interaction and nature of mutation-dependent dysregulation remains unresolved (*26*) and microglia have never been studied. A range of studies suggest interaction of C9ORF72 at several points in the autophagy pathway. These include, p62 dependent recruitment of cargo, followed by the trafficking of ULK-1 autophagy initiation complex for autophagosome formation through its interaction with Rab proteins, and, is also known to be involved in later stages of autophagy during the fusion of autophagosome with the lysosome (*36*). Now, at the cellular level, C9ORF72 forms a stable multi-functional complex with SMCR8 and WDR41 to execute these autophagic and vesicle trafficking events (*37, 38*). This was further supported by the independent and combinatorial knock down of C9ORF72 and SMCR8 revealing their role in the initiation of autophagy, however they may have opposing roles in the regulation autophagy flux (*26*). Therefore, further studies will be required to understand the precise role of C9ORF72 in regulating autophagy in immune cells.

Notwithstanding these mechanistic insights, autophagy and the C9ORF72 interactome has largely been studied from a neurocentric perspective (*26*). However, the role of C9ORF72 on microglial autophagy and the impact of mutation-dependent dysregulation on microglial biology and function is unclear. Our study emphasizes the value of studying cell-type specific behaviour to not only uncover novel interacting partners but also address non-neuronal consequences of dysregulated autophagy. We report a novel interaction of C9ORF72 with UFM 1 (Ubiquitin fold modifier 1) and UFL1 (an UFM1 E3 ligase), which is essential for haematopoietic stem cell function as well as the transcription factor PU.1 that collectively suggests additional roles for C9ORF72 in myeloid cell function (*39*). Interestingly, UFMylation has been recently shown to be involved in ER-phagy, thereby providing indirect evidence on the role of C9ORF72 in UFMylation-induced ER-phagy (*40*). Furthermore, elevated IFN response to proinflammatory stimuli such as LPS is observed in UFMylation-deficient cells, similar to the finding in C9-/-mice, suggestive of a mutually dependant mechanistic outcome (*13, 41*). The finding that C9ORF72 in human iPSC-derived microglia-like cells is a binding partner of key mediators of the autophagy initiation complex such as SMCR8 and WDR41 is consistent with previous reports (*26, 42*). Interestingly, rodent studies from SMCR8-/- and WDR-/-mice also exhibited an exaggerated immune response similar to our findings of human mC9orf72 myeloid cells upon LPS stimulation. This deficit was reversed by knockout of endosomal TLRs consistent with SMCR8 negatively regulating TLR signalling. Together this suggests that functional loss of the C9ORF72-SMCR8-WDR41 complex drives an altered immune state through reduced autophagy (*43*). However, TLRs are not the only immune responders reliant on endolysosomes for their activity, other inflammatory mediators such as NLRP3 and members of NFκb pathway are also selectively degraded by autophagy (*44*). The NLRP3 inflammasome is a cytosolic multi-protein complex, which responds to cellular stress/endogenous stimuli like secreted ATP and reactive oxygen species and promotes the activation of interleukin-1β (IL-1β) and IL-18 through active caspase-1 (*45, 46*). Furthermore, Nfκb upon exposure to inflammatory stimuli is activated through signalling cascades including the IKK kinase complex resulting in nuclear translocation of NF-κB to initiate the transcription of pro-inflammatory cytokines (*47*). Indeed, excitotoxicity, a key pathogenesis in ALS, is also reported to elicit microglial activation; however, it was not known if the enhanced vulnerability of C9 motor neurons to excitotoxicity could contribute to microglial activation in *C9orf72*-ALS and whether such activation plays a causal role in motor neuronal death. Interestingly, cytokines are already known to elicit neuronal death via nitric oxide and free radicals(*48*). Given that autophagy regulates (i) NLRP3 inflammasome complex and (ii) IKKα/β kinases that result in activation of the Nfκb pathway (*49*), our pulse-chase experiments using LPS on microglial monoculture and AMPA (excitotoxic stimulus) on co-culture of MN-MG revealed persistent activation of NLRP3 inflammasome and NF-κB signalling in mC9-MG, leading to concomitant increased production of cytokines in absence of autophagy mediated feedback inhibition. This result demonstrated for the first time that cell intrinsic vulnerability of mC9-MGs to triggering stimuli contributes to enhanced motor neuronal death in *C9orf72*-ALS. Indeed, our finding of increased NLRP3 activation and reduced autophagy activation in human hiPSC-MG is in line with the existing literature reporting elevated levels of NLRP3 in astrocytes in sporadic ALS cases and suboptimal autophagy in the ventral horn of ALS patients (*46, 50, 51*). Pharmacological correction of this deficit using rapamycin raises the possibility of future therapeutic intervention, noting also the increasing identification of a range of drugs that appear to boost autophagy of relevance to multiple neurodegenerative diseases (*52–55*).

Emerging evidence suggests a cross-talk between autophagy and phagocytosis. This includes recent findings that show that LC3-associated phagocytosis (LAP) in myeloid cells is an efficient intracellular eliminator of engulfed extracellular cargo, including misfolded proteins, (*56, 57*). Our finding of reduced phagocytosis in m*C9orf72* myeloid cells that is ameliorated by boosting autophagy is consistent with these reports. The relevance of these findings to the TDP-43 accumulation in *C9orf72*-ALS would be of great interest to investigate in respect to microglial pathology as well as potential TDP43 spread given the report that microglial depletion appears to facilitate axonal spread of TDP-43 (*58*). Further evidence suggesting an important role for LC3 is the finding that microglia-specific LC3-associated endocytosis (LANDO)-deficient rodent models exhibited widespread tau pathology and neuronal death (*59*).

The absence of reliable and logistically simple-to-obtain biomarkers for disease prediction and progression remains a challenge for developing and delivering individualised therapies. Our finding of a quantitative reduction in C9ORF72 in PBMCs from both carriers and symptomatic patients complements emerging assays of DPRs from CSF and / or PBMCs (*60*). Furthermore, the correlation of reduced abundance with functional deficits potentially opens new approaches to the use of C9ORF72 levels along with functional assays as an exploratory biomarker for monitoring disease progression and enabling precision medicine.

While our findings are consistent with a loss-of-function model, this is not mutually exclusive with a double model of loss- and gain-of-function. Such a two-hit model is supported by transgenic and *in vitro* human neuronal studies, showing that reduced levels of C9ORF72 lead to impaired autophagy and dipeptide repeat protein accumulation (*14, 24*).

In conclusion, our study uncovers the impact of impaired autophagy on m*C9orf72* human microglial function, resulting in sustained activation of the NLRP3 inflammasome and NF-κB signalling pathways, which led to an enhanced motor neuronal death following excitotoxic insult. Together, this suggests a state of enhanced vulnerability of m*C9orf72* microglia to triggering stimuli and cellular stress. Exogenous pharmacological activation/boosting of autophagy with mTOR inhibitors ameliorated this pro-inflammatory phenotype, thus providing a target for translational research.

## Materials and Methods

### Differentiation and characterisation of microglia (hiPSC-MG) from human iPSCs

A control iPSC line (CTRL), two *C9orf72* ALS/FTD patient iPSC lines (mC9-1 and mC9-2) harbouring the GGGGCC repeat expansion and their respective isogenic control lines (isoC9-1 and isoC9-2) used in this study were generated with full Ethical/Institutional Review Board approval at the University of Edinburgh, as previously reported by our group (*11, 32*). iPSCs were maintained on a monolayer of irradiated mouse embryonic fibroblast (Thermo Fisher Scientific) in Essential 8 Medium (E8, Gibco) in plates coated with Matrigel Growth Factor Reduced Basement Membrane Matrix and were passaged at 80% confluence with 1mg/ml collagenase/0.5mg/ml dispase (Thermo Fisher Scientific). All iPSCs were confirmed to be sterile, mycoplasma-free and G-banding was performed at regular intervals to rule out any acquired chromosomal abnormality in the iPSCs.

Human microglia-like-cells (hiPSC-MG) were generated following a protocol previously described by our group (*23*). Briefly, myeloid progenitors expressing CX3CR1, CD11b and CD45 were derived from mesodermally-patterned embryoid bodies cultured in myeloid precursor media containing 100ng/ml M-CSF (Peprotech) and 25ng/ml IL-3 (Gibco) for 4-6 weeks. These myeloid precursors were then differentiated to microglia-like-cells (hiPSC-MG) on poly-D-lysine treated gelatinised plates (0.01%) in the presence of 100ng/ml IL-34 (Peprotech), 10ng/ml GM-CSF (Peprotech) and line matched neural precursor cell conditioned media (NPC-CM), which was supplemented in an increasing gradient (10%-50%) every other day. The hiPSC-MG obtained from control (ctrl), C9orf72 mutant (mC9) and isogenic control (isoC9) were immune-stained with microglial signature markers, such as TMEM119, P2Y12 and PU.1. The catalogue codes for the antibodies alongside working concentration and reagents are mentioned in resource table.

### Immunocytochemistry

hiPSC-MG grown on coverslips were fixed in 4% paraformaldehyde (PFA; Agar scientific) in phosphate-buffered saline (1X PBS) for 10 min at room temperature. After fixation, cells were washed three times with 1X PBS, permeabilised with 0.1% Triton-X100 (Thermo Fisher Scientific) in 1X PBS for 5 min and blocked with 3% BSA (Europa Bioproducts) in 1X PBS for 1hr at room temperature. Incubation with primary antibodies for two hours was followed by addition of appropriate secondary antibodies. The cells were then washed three times with 1X PBS, counterstained with 4′,6-diamidino-2-phenylindole (DAPI; Thermo Fisher Scientific) and mounted on to glass slides in FluorSave™(Millipore). The slides were observed in Zeiss LSM confocal microscope. The catalogue codes for the antibodies and reagents are mentioned in resource table.

### Flow cytometry

The myeloid precursors were collected from the supernatant, centrifuged, washed with 1X PBS and were treated with 1% Fc block (Miltenyi Biotec) in 1X PBS for 10 min at room temperature, before incubating them with primary antibodies for one hour on ice. Samples were then washed twice with 1X PBS and transferred onto round-bottom polystyrene tubes (BD Falcon) until assessment. The stained cell samples were analysed using a FACS LSR Fortessa (4 lasers) flow cytometer (BD Biosciences), and the data were processed using the FlowJo software.

### High throughput imaging and analysis platform for the quantification of microglial phagocytosis

Microglial differentiation was performed in a 96 well black/clear bottom plate with optical polymer base (Thermo Scientific™) for 12 days. pHrodo-conjugated zymosan beads (Thermo Fisher Scientific) were added to the hiPSC-MGs at a concentration of 0.5 mg/ml and the plate was immediately loaded into ImageXpress Micro 4 High-Content Imaging System, a high throughput live imaging system maintained at 37 degrees and 5% CO2 to monitor a real-time uptake of zymosan beads by the hiPSC-MGs. Prior to loading, hiPSC-MG were co-stained with Hoechst 33342 (Thermo Fisher Scientific). The cells were imaged using a 20X objective at an interval of 15 min for 2.15 hours (135 min) and 4 field of view were captured per well. All lines reported in this study had minimum of 3 biological replicates and 6 wells from each replicate per line were examined. Thus 24 field of view per replicate from each line were analysed for the zymosan bead uptake assay.

In order to determine the phagocytic index of individual genotypes (mutant and isogenic control) images were analyzed using Definiens Developer XD. Nuclei were first detected using a fluorescence threshold to separate background pixels from Hoechst-stained objects. Any holes in objects were filled and the objects were smoothened by shrinking and growing each object by 2 pixels. Objects were then classified as nuclei based on their size (excluding small and large objects) and their elliptical fit (only objects with an elliptical fit greater than 0.8 were classified as nuclei). The nuclei were then expanded by 50 pixels and within this area we used a contrast threshold to identify the borders of pHrodo objects. pHrodo objects were smoothened by shrinking and growing each object by 2 pixels and then filtered using a fluorescence threshold to remove any objects that were not significantly brighter than the background (>2- fold change). From each image we calculated the number of nuclei (Hoechst-stained objects), the total number of spots (pHrodo beads), and the data was plotted as beads/cell after normalising the total number of spots with total number of nuclei.

### LPS induction and assessment of NLRP3 and NF-κB activation

hiPSC-MGs were stimulated with 100 ng/ml lipopolysaccharide (LPS; Sigma-Aldrich) for 8 hr and followed by a washout. Cell lysates were collected at 4h, 8h and 12h post washout respectively to examine the dynamics of NLRP3 and p62 by western blotting. For testing the dynamics of NF-κB signalling by immunochemistry, cells post LPS washout were fixed at 8h and 12h respectively and stained with anti-NF-κB (Abcam) and anti LC3B (Enzo Lifesciences) and observed in Zeiss LSM confocal microscope. The details of the dilution for individual antibodies are mentioned in Resource table.

For assessing NF-κb activation, the Carl Zeiss image files were imported into Imaris version 8.0.2, the nuclear region was identified using DAPI staining and the intensity of the nuclear NF-κb was measured in voxels using Imaris software. Results were exported from Imaris in CSV formats and graphs represent the colocalised voxels of DAPI and nuclear NF-κB.

### Cytokine ELISA

The concentration of IL-6 and IL-1β in cell supernatants was measured using DuoSet ELISA Development Systems (R&D Systems). hiPSC-MGs generated from control iPSC (ctrl), *C9orf72* mutant iPSC (mC9-1 and mC9-2), their respective isogenic (isoC9-1 and isoC9-2) in a 12-well dish were treated with LPS or vehicle control (1XPBS) for 8h and following a 12h washout, the cell-free supernatants were collected from the cultures after a brief centrifugation at 2000 RPM for 5 min. The samples were then assessed in triplicate for IL-6 and IL-1β and the concentration (pg/ml) obtained was further normalised to the 100ug of protein content (pg/ml) in each condition for monocultures of microglia and 500ng of protein for MN-MG co-cultures. The catalogue codes for the ELISA kits are mentioned in resource table.

### Co-culture of spinal motor neuron (MN) and microglia (MG) and quantification of neuronal number and microglial number in the MN-MG co-culture

Enriched spinal motor neuron culture was generated from *C9orf72* patient and isogenic iPSC lines using an established protocol (*11, 32*). Briefly, iPSCs were lifted using Accutase and were neuralised by dual SMAD inhibition using 20μM SB431542, 0.1μM LDN-193189 and 3μM CHIR-99021). The spheres were maintained as a suspension culture for 2 weeks. Post two weeks, the spheres were dissociated using 0.05% trypsin-EDTA and 30,000 cells were plated on 96 well plates and on coverslips in 24-well plates. These cells were cultured in motor neuron neurotrophic factor (MN-NF) media for 3 weeks. Microglial cells were differentiated from the myeloid precursors and were lifted using Accutase for co-culturing with motor neurons. Post two weeks, 5,000 microglial cells (hiPSC-MG) were added on the motor neurons and were co-cultured for one week, before challenging them with 100 μM AMPA for 24 hours.

Following AMPA addition, the cells were fixed with 4% PFA and stained with β-III tubulin, Iba-1, p62, NLRP3, nfκb for further experiments and quantification. DAPI, beta-III Tubulin and IBA-1 as above were imaged using an ImageXpress micro-confocal with a 20X plan Apo objective and 9 fields of view per well. All lines reported in this study had minimum of 3 biological replicates. A custom analysis module was designed with MetaXpress software to count the number of beta-III Tubulin+ neurons and the number of IBA-1+ microglia based on their size and intensity above background. The catalogue codes for the reagents and antibodies are mentioned in resource table.

### Western blot analysis

Western blot analysis was carried out according to established protocol (*61*). Briefly, the cells were lysed in lysis buffer (20 mM Tris-Cl pH 7.4, 1 mM EDTA, 150 mM NaCl, 1% Triton X-100, 1 mM EGTA,, 1 mM phenylmethylsulfonyl fluoride). The protein concentration was calculated using the BCA Protein Assay Reagent (23225, Pierce™ Thermo Fisher Scientific). Protein samples were separated by SDS-PAGE (Thermo Fisher Scientific) and transferred to polyvinylidene difluoride (PVDF) membranes (Millipore, Bedford, MA, USA). The membranes were blocked with 5% skimmed milk and incubated at 4 °C overnight with primary antibodies followed by addition of secondary antibody conjugated with horseradish peroxidase. The protein band signals were detected by ECL chemiluminescence substrate (Amersham) and quantified by Image J software. The catalogue codes for the reagents and antibodies are mentioned in resource table

### Autophagy flux assay using mRFP-GFP-LC3 dual reporter probe in hiPSC-MG

Monocultures of hiPSC-MGs were lifted with 1X Accutase (Merck) and transduced with lentivirus expressing mCherry-EGFP-p62 at a MOI (multiplicity of infection) of 100 to obtained a transduction efficiency of 80%. The transduced live cells were visualised and fluorescence images were captured after 48h using Zeiss 710 confocal microscope, followed by the manual counting of GFP and RFP puncta per cell employing Image J software. Colocalised puncta for GFP and RFP are indicated as autophagosomes (as represented in yellow bar in Fig 3D) and red puncta are considered as autolysosomes (as represented in red bar in Fig 3D).

### Generation of C9 knock-out iPSC line

C9ORF72 KO was generated using the following strategy, two small gRNAs were used to delete a 100 bp fragment in exon 2 of C9ORF72 gene. Bi-allelic deletion and following non-homologous end joining event resulted in-frame STOP codon, thus, leading to premature termination of C9ORF72 protein translation. Experimentally, two sgRNAs (gRNA-1 5’-CAACAGCTGGAGATGGCGGT-3’, gRNA-2: 5’-ATTCTTGGTCCTAGAGTA-3’) were cloned individually in pSpCas9(BB)-2A-Puro (PX459) as per (*62*). Control iPSCs (CNTR-1, 8x 10^5^) at 70-8-% confluence were dissociated into single cells using 1x Accutase (Merck) and were nucleofected with 2 μg Cas9-gRNA-1, 2 μg Cas9-gRNA-2, and 1 μg pEGFP-Puro (for positive selection of transfected cells), using the P3 Primary Cell 4D-Nucleofector TM X Kit (Lonza), program CA-137) on a Lonza 4D-Nucleofector™ X Unit (Lonza). Transfected iPSCs were plated on to Matrigel coated dish and cultured using E8 media (Gibco) and ROCK inhibitor (10 μM)(Tocris). Selection with 1 μg/ml puromycin (ThermoFisher) was commenced 24h post-nucleofection and continued for 24h. Cells were dissociated and plated at a low density (1×10^4^ cells/10 cm plate) for clonal selection upon confluence. Ninety-six individual clones were manually picked and screened for 100 bp deletion in exon 2 using following primers C9-KO_Fw: 5-ACAGGATTCCACATCTTTGACT-3’ and C9-KO_Rev: 5’-GCGATCCCCATTCCAGTTTC-3’. The positive C9-KO iPSC clone was confirmed by Sanger sequencing (Suppl. Fig 3A) and immunoblot analysis (Suppl. Fig 3A), which validated the loss of C9ORF72 protein.

### Generation of EGFP-C9ORF72 knock-in iPSC line

sgRNAs were designed using the CRISPR design toolkit (http://crispr.mit.edu) and cloned into pSpCas9(BB)-2A-Puro (PX459) V2.0 plasmid (Addgene: #62988) (*62*). sgRNAs were designed using CRISPR toolkit (http://crispr.mit.edu). Sequences of gRNAs are as follows: gRNA(1): 5’-GCATTTGGATAATGTGACAGT-3’; gRNA(2): 5’-GTCACATTATCCAAATGCTC-3’; gRNA(3): 5’-GACCGCCATCTCCAGCTGTTG-3’. Genomic targeting efficiency was determined in the Ctrl iPSC line via T7 Endonuclease I assay PCR amplicon flanking the C9orf72 target site (generated using T7_Fwd and T7_Rev primers as listed in Resource table) was denatured, re-annealed and subsequently digested. All three sgRNAs were effective in generating a double strand break, and gRNA(1) was taken forwards because it introduces a double strand break closer to the end of the 5’UTR of the *C9orf72* gene locus.

A targeting vector comprising of a sequence coding for an enhanced GFP (EGFP) protein, flanked by ~1 kb homology arms (HA) were cloned into pMTL23 vector backbone using Gibson assembly (*63*). Homology arms and EGFP incorporating a Kozak consensus sequence were generated through PCR amplification, using primers listed in Resource Table. The pAAV-FLEX-EGFPL10a plasmid (Addgene) (*64*)was used to generate the EGFP fragment. Sequences of these fragments are listed in Supplementary table 1. They were assembled by NEBuilder HiFi DNA Assembly Cloning Kit (NEB).

Ctrl iPSCs at 70-80% confluence were dissociated into a single-cell suspension using 1X Accutase (Merck). 8×105 cells were nucleofected with 2 μg of Cas9-gRNA1 plasmid, 2 μg of targeting vector and 1 μg of puromycin resistance vector (for positive selection of transfected cells) using the Lonza 4D-Nucleofector X unit (program CA137) following the manufacturer’s instructions. Transfected cells were plated onto Matrigel (BD)-coated dishes in the presence of E8 medium supplemented with ROCK inhibitor (10 μM; Y-27632, Tocris). 24 hours post nucleofection, cells were subjected to 24 hours puromycin selection (1 μg/ml). Upon confluence, cells were dissociated again into single cells and plated down at low density (2000 cells/10 cm dish) for clonal selection. After 8-10 days, single colonies were isolated and transferred to Matrigel (BD)-coated 96-well plates. 300 clones were picked manually. Locus-specific PCR was performed using C9Ext_Fwd (located in intron 1 of C9orf72, external to the 5’ homology arm) and C9_Rev (located in the 3’ homology arm within intron 2-3). The sequence of the primers is listed in resource table).

### Co-immunoprecipitation of EGFP-C9ORF72 and mass spectrometry

EGFP-C9ORF72 hiPSC-MGs were grown on 10-cm dish and were lysed using a 20 mM HEPES pH 7.5, 150mM NaCl, 5 mM MgCl2, 10% glycerol, 0.5% NP-40, 10 mM sodium glycerophosphate, 10 mM sodium pyrophosphate, 0.1 μM microcystin-LR, 1 mM sodium orthovanadate, 100 nM GTPgS, Complete EDTA-free protease inhibitor cocktail. 350 μg of protein from ctrl-MG and EGFP-C9ORF72-MG (two biological replicates, 4 technical replicates) were subjected to immunoprecipitation using 10 μl of Chromotek GFP-Trap beads. Samples were incubated on an end-to-end rotator in cold room (4°C) for 90 minutes followed by the washes with lysis buffer. On-bead tryptic digestion followed by 8-plex TMT labelling was performed as described earlier (*65*). TMT labelling was verified by taking 5μl from each sample was pooled and analysed by LC-MS/MS analysis. TMT labelling efficiency was found to be >98%, remaining samples were quenched by adding 5μl of 5% Hydroxyl amine. Samples were pooled and vacuum dried. To increase the depth the pooled TMT labelled sample was fractionated using mini high-pH RPLC strategy on homemade C18 stage-tips (*65*), eluted peptides were vacuum dried until LC-MS/MS analysis.

### LC-MS/MS analysis

The basic reverse-phase liquid chromatography (bRPLC) fractions were reconstituted in 15 μl of 0.1% formic acid and 3% ACN buffer and subjected to LC-MS/MS/MS analysis on Orbitrap Exploris 480 hybrid mass spectrometer that is interfaced with 3000 RSLC nano liquid chromatography system. Sample was loaded on to a 2 cm trapping column (PepMap C18 100A – 300μm. Part number: 160454. Thermo Fisher Scientific) at 5 ul/min flow rate using loading pump and analyzed on a 50cm analytical column (EASY-Spray column, 50 cm × 75 μm ID, Part number: ES803) at 300 nl/min flow rate that is interfaced to the mass spectrometer using Easy nLC source and electro sprayed directly into the mass spectrometer. LC gradient was applied from a 3% to 25% of solvent-B at 300 nl/min flow rate (Solvent-B: 80% CAN) for 100 minutes and increased to 45% solvent-B for 10 minutes and 40% to 99% Solvent-B for 5 minutes which is maintained at 90% B for 10 minutes and washed the column at 3% solvent-B for another 10 minutes comprising a total of 145 min run with a 120-minute gradient in a data dependent MS2 mode. The full scan MS1 was acquired at a resolution of 120,000 at m/z 200 between 350-1200 m/z and measured using ultra-high filed Orbitrap mass analyzer. Precursor fit threshold of 70% at 0.7 Da isolation width filter enabled accurate isolation of precursor isotope envelope for the MS2 fragmentation. The top 10 precursor ions were targeted which are isolated using Quadrupole mass filter at 0.7 Da isolation width for the MS2 and fragmented using 36% higher-energy collisional dissociation (HCD) analyzed using Ultra high-filed Orbitrap mass analyzer at a resolution of 45,000 at m/z 200. AGC target for MS1 and MS2 were set at 300% and 100% respectively with a maximum ion injection times of 28 ms for MS1 and 110 ms for MS2.

### Mass spectrometry data analysis and bioinformatics analysis

The MaxQuant software suite (*66*) version 1.6.10.0 was used for database search with the following parameter. Reporter ion MS2 type: 8plex TMT with PIF (Precursor intensity factor) of 0.7 to have accurate reporter ion intensities. The TMT isotopic reporter ion correction factors were manually entered as provided by the manufacturer. The following group specific parameters were used, A built-in Andromeda search engine was used by specifying Trypsin/P as a specific enzyme by allowing 2 missed cleavages, minimum length of 7 amino acids, Oxidation of (M), Acetyl (Protein-N-terminal), Deamidation N and Q were selected as variable modifications. Carbamidomethylation Cys was selected as fixed modification. First search tolerance of 20 ppm and main search tolerance of 4.5 ppm were selected. Global Parameters: Uniprot Human protein database (release 2017-02; 42,101 sequences) was used for the database search and 1% peptide and protein level FDR was applied. For protein quantification, min ratio count was set at 2 for accurate reporter ion quantification. The MaxQuant output protein group text files were processed using Perseus software suite (*67*), version 1.6.10.45 was used. The data was filtered for any proteins that are flagged as common contaminants and reverse hits. In addition a common background binders from Crapome database (*68*) were filtered and finally minimum two unique peptide were retained for the downstream analysis. The TMT reporter ion intensities were log2 transformed and subsequently the TMT reporter tags of Control and GFP-C9orf72 conditions were categorized to perform statistical analysis. Two sample Welch’s T-test was performed by applying 1% and 5% permutation-based FDR to identify the differentially enriched and significant protein groups between GFP-C9orf72 and control groups.

### *C9orf72* case and age- and sex-matched control cohort for PBMC collection

All clinical meta-data were collected as part of Scottish Motor Neurone Disease Register (SMNDR) and Care Audit Research and Evaluation for Motor Neurone Disease (CARE-MND) platform (ethics approval from Scotland A Research Ethics Committee 10/MRE00/78 and 15/SS/0216). All cases had corresponding whole-genome sequencing and diagnostic repeat prime PCR, demonstrating pathogenic repeat lengths in the C9orf72 locus (*69*). Blood samples from cases and age- and sex-matched healthy volunteers were taken under Lothian NRS Human Annotated Bioresource ethics (15/ES/0094), in line with the Human Tissue (Scotland) Act 2006, after obtaining signed informed consent, as part of the Scottish Regenerative Neurology Tissue Bank.

### PBMC collection and macrophage differentiation

Venepuncture was performed and ~16 ml of blood collected in two BD Vacutainer® CPT™ Mononuclear Cell Preparation Sodium Citrate Tubes at the Anne Rowling Regenerative Neurology Clinic, University of Edinburgh. Samples were gently inverted ~5 times and were immediately centrifuged for 20 min at 1600 g at room temperature, permitting the formation of a physical barrier between the mononuclear cells in plasma and the erythrocytes and granulocytes. Mononuclear cells were collected from the buffy coat as described in (*70*) and were differentiated to functional macrophages using 80ng/ml MCSF in RPMI media (Lonza) for 7 days. These macrophages were then assayed for phagocytosis using zymosan beads (Thermo Fisher Scientific) and cytokine production (IL6 and IL-1β) was measured across unstimulated and LPS stimulated condition employing ELISA (R&D Systems).

### Statistical analysis

Statistical analysis was performed using Prism version 8.4.0 (GraphPad Software) and differences were considered significant when p <0.05 (*p < 0.05, **p < 0.01, ***p < 0.001. For comparison of normally distributed data of two groups, two-tailed unpaired Student’s t– test was performed. Comparisons of data consisting of more than two groups where they vary in two factors, was performed using two-way ANOVA and subsequent Tukey’s and Sidak’s multiple comparisons test. Detailed statistical information for each figure including statistical tests performed are elaborated in the respective figure legends.

## Acknowledgements

The authors thank Dr. Daniel Scott, University of Nottingham and Dr. Robert Layfield, University of Nottingham, for sharing the mCherry-GFP-p62 plasmid. The authors also thank Dr. Pamela Brown, University of Edinburgh, for generating the lentiviral vectors.

## CRediT Contribution

Poulomi Banerjee (PB): Conceptualization, Validation, Investigation, Visualization, Methodology, Writing — original draft, Project administration, Writing – review and editing Arpan R Mehta (ARM): Methodology, Investigation, Resources, Writing – review and editing

Raja S Nirujogi (RSN): Methodology, Investigation, Writing - Review & Editing James Cooper (JC): Methodology, Resources

Owen G James (OGJ): Methodology, Software, Formal analysis Jyoti Nanda (JN): Investigation

James Longden (JL): Methodology, Software, Formal analysis Karen Burr (KB): Resources

Andrea Salinzger (AS): Methodology, Investigation Evdokia Paza (EP): Investigation

Judith Newton (JN): Resources (Clinical work)

David Story (DS): Resources (administrative support)

Suvankar Pal (SP): Resources (Recruitment of patients via ARRNC)

Colin Smith (CS): Resources (post-mortem samples), Supervision, Writing - Review & Editing

Dario R Alessi (DRA): Methodology, Validation, Supervision Investigation, Writing - Review & Editing

Bhuvaneish T Selvaraj (BTS): Conceptualisation, Methodology, Writing - Review & Editing, Project administration.

Josef Priller (JP): Conceptualization, Supervision, Project administration, Visualization, Writing - Review & Editing

Siddharthan Chandran (SC): Conceptualization, Resources, Supervision, Project administration, Visualization, Writing - Original Draft, Writing - Review & Editing, Funding acquisition

