## Supplementary material for "Cell-autonomous immune dysfunction driven by disrupted autophagy in *C9orf72*-ALS iPSC-derived microglia contributes to neurodegeneration": Suppl Figs

**This PDF file includes:**

Supplementary figs 1-11

**Supplementary Fig 1 (Supp. To Fig.1)**

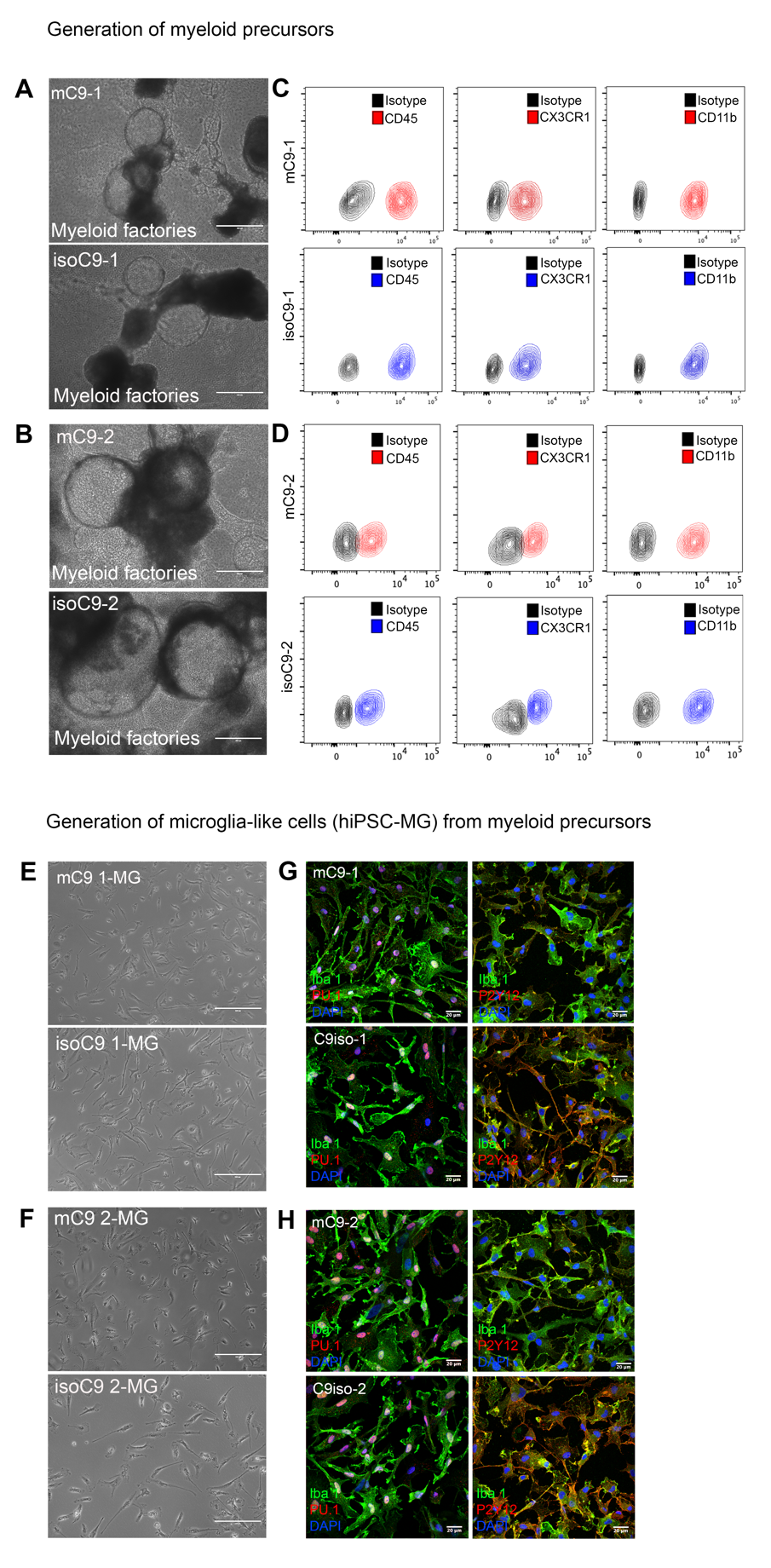

**Supplementary Fig 1- Generation and characterisation of human microglia-like cells(hiPSC-MG) from mC9 and isoC9 iPSCs:** **(A, B)** Phase contrast images showing the formation of myeloid factories from two pairs of C9 mutant (m*C9*-1 and m*C9*-2) and their respective isogenic iPSC lines (iso*C9*-1 and iso*C9*-2), Scale bar=200μm **(C,D)** Myeloid precursors produced by the mC9 and isoC9 myeloid factories show comparable profile for key myeloid precursor markers- CD45,CX3CR1 and CD11b (**E,F)** Myeloid precursors generated from the myeloid factories were differentiated into human microglia-like cells (hiPSC-MG) by treating them with IL34 ,GM-CSF supplemented with neural precursor cell conditioned media (NPC-CM) **(G,H)** Representative immuno-staining of two pairs mC9-MG and isoC9-MG showing staining for Iba-1, purinergic receptor P_2_Y_12_ and myeloid marker PU.1; scale bar=20μm. Data representative of n=3.

**Supplementary Fig 2 (Suppl. To Fig 2)**

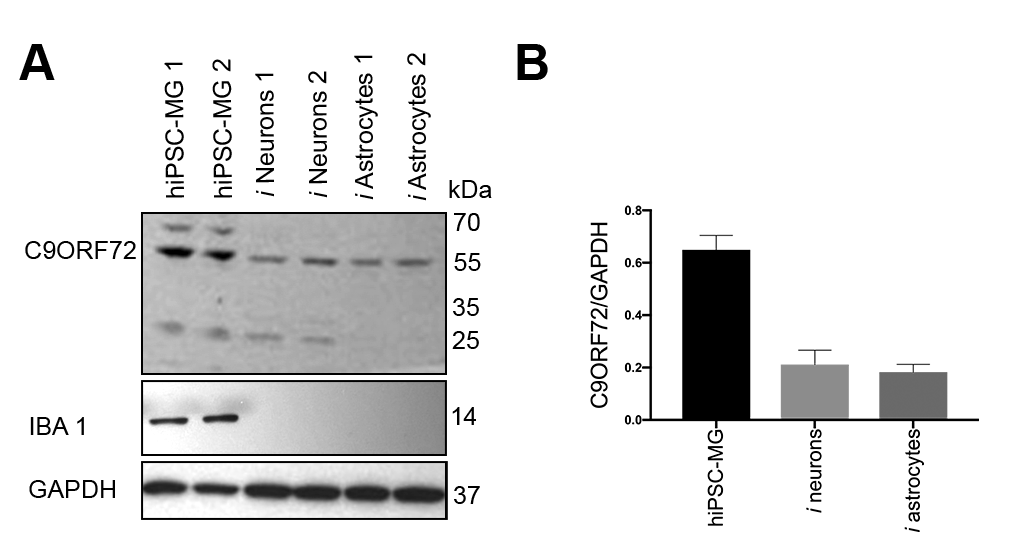

**Supplementary Fig 2- Comparison of C9ORF72 level across iPSC derived microglia (hiPSC-MG), neurons *(ipsc* neurons) and astrocytes (*ipsc* astrocytes)**: **(A)** Immunoblots showing the higher abundance of C9ORF72 in hiPSC derived microglia (hiPSC-MG) when compared to hiPSC derived neurons (*i* neurons) and astrocytes (*i* astrocytes) generated from two independent iPSC lines, immunoblot for IBA-1 confirms the identity of hiPSC-MG **(B)** bar graphs show the densitometric quantification of C9ORF72 levels over GAPDH for different cell types derived from iPSC, data represent mean +/-SD across 3 biological replicates.

**Supplementary Fig 3 (Suppl. To Fig 2)**

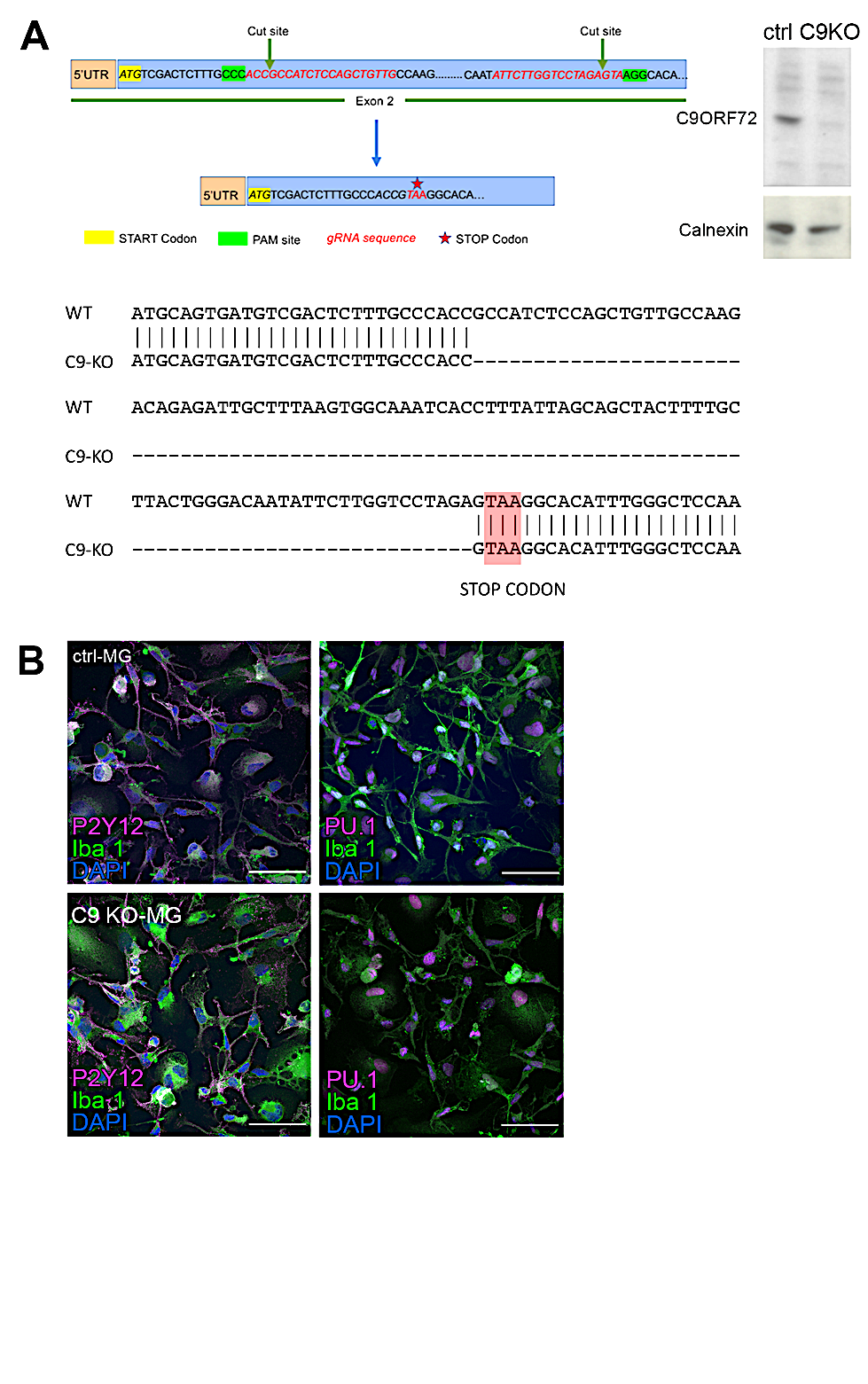

**Supplementary Fig 3**- **Generation and characterisation of C9KO-MG from C9KO iPSC line:** **(A)** Strategy for the generation of C9 knock out iPSC-line, two gRNAs were used to delete a 100 bp fragment in exon 2, and following non-homologous end joining event, this resulted in the insertion of an in-frame STOP codon, immunoblot (right) confirming the absence of C9ORF72 protein in the C9KO iPSC line, Sanger sequencing depicting the deletion of 100bp region in exon2. Highlighted region shows the in frame stop codon **(B)** Represents the immuno staining of C9KO iPSC derived microglia (C9KO-MG) and ctrl-MG from the same genetic background confirming the positive staining for microglial markers such as P_2_Y_12_ and PU.1

**Supplementary Figure: 4 (Suppl. to Fig 2)**

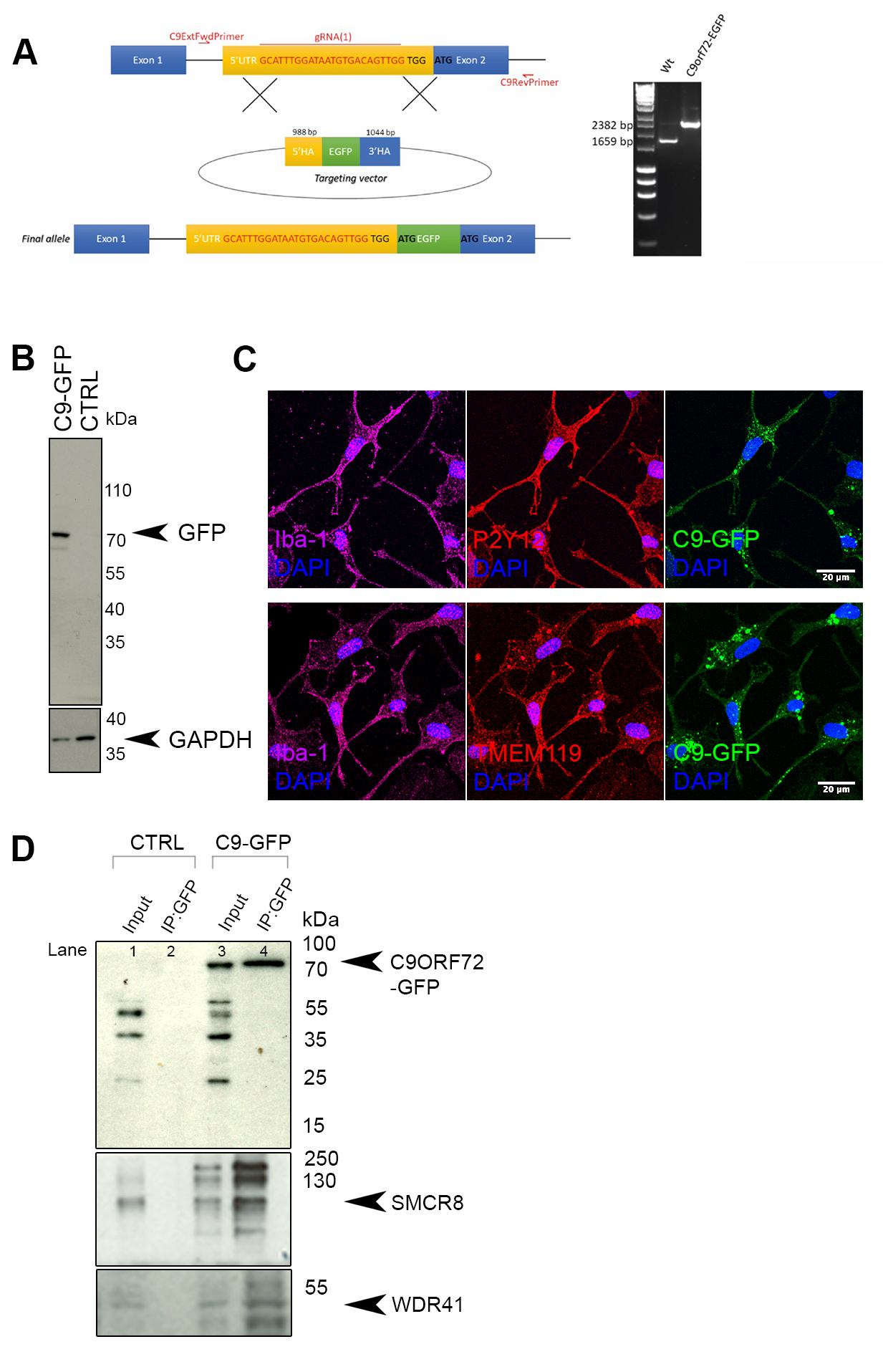

**Supplementary Fig 4- Generation and characterisation of C9GFP-MG from *C9orf72*-GFP iPSC line and validation of the C9ORF72 interactome**

**(A)**Schematic diagram (left) depicting gene targeting strategy, and location of guide RNA and forward and reverse locus-specific primers (**see Supplementary Table 1**) for primer pair sequence) used as part of the validation process for the homozygous clone. Homology arms (HAs) as shown: 5’HA comprises part of intron 1 and the 5’UTR of exon 1; 3’HA comprises exon 2 and part of intron 2. Agarose gel electrophoresis (right) showing genotyping screen using locus-specific primers. Agarose gel electrophoresis shows a single higher band for the homozygous EGFP-C9orf72 clone (2382 bp) when compared with the wild-type band (1659 bp). **(B)** Immunoblot demonstrating the expression of GFP-C9 fusion protein in C9GFP-MG (corresponding to 70kDa) as opposed to no bands in ctrl-MG **(C)** Representative images of immunofluorescence staining for P_2_Y_12_ and TMEM119 confirming the identity of C9GFP-MG **(D)** Western blotting for GFP-C9ORF72 fusion protein (corresponding ~70kDa in C9GFP -MG and 55kDa in ctrl-MG), SMCR8 (105kDa) and WDR41 (48.5kDa) in GFP pull down samples of C9GFP-MG and ctrl-MG confirming the interaction of C9ORF72 with SMCR8 and WDR41.

**Supplementary Fig 5 (Suppl. To Fig 3)**

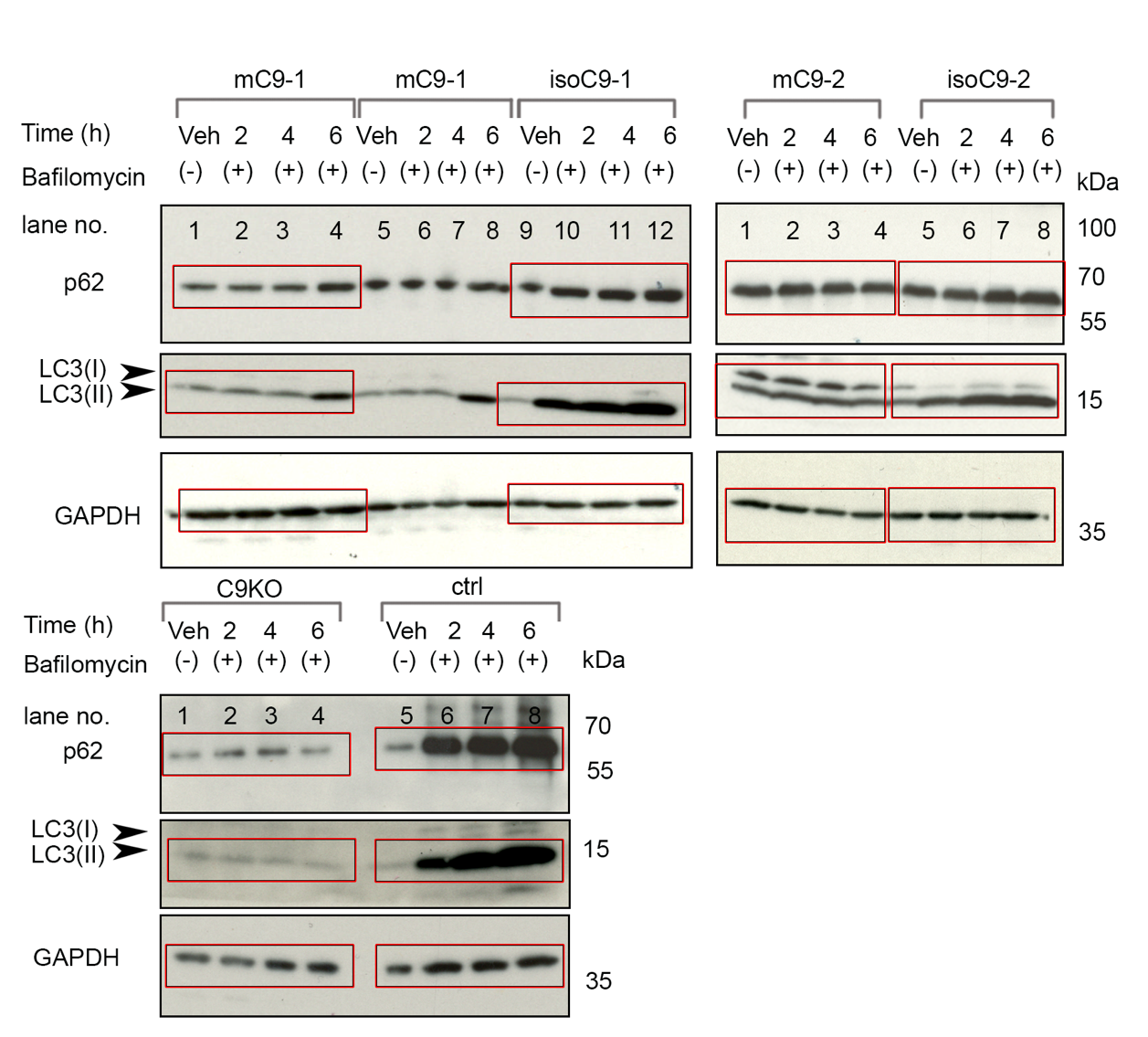

**Supplementary Fig 5: Full blot of Fig 3 B:** The bands highlighted in red rectangle have been considered for the main figure (Fig 3B)

**Supplementary Fig :6**

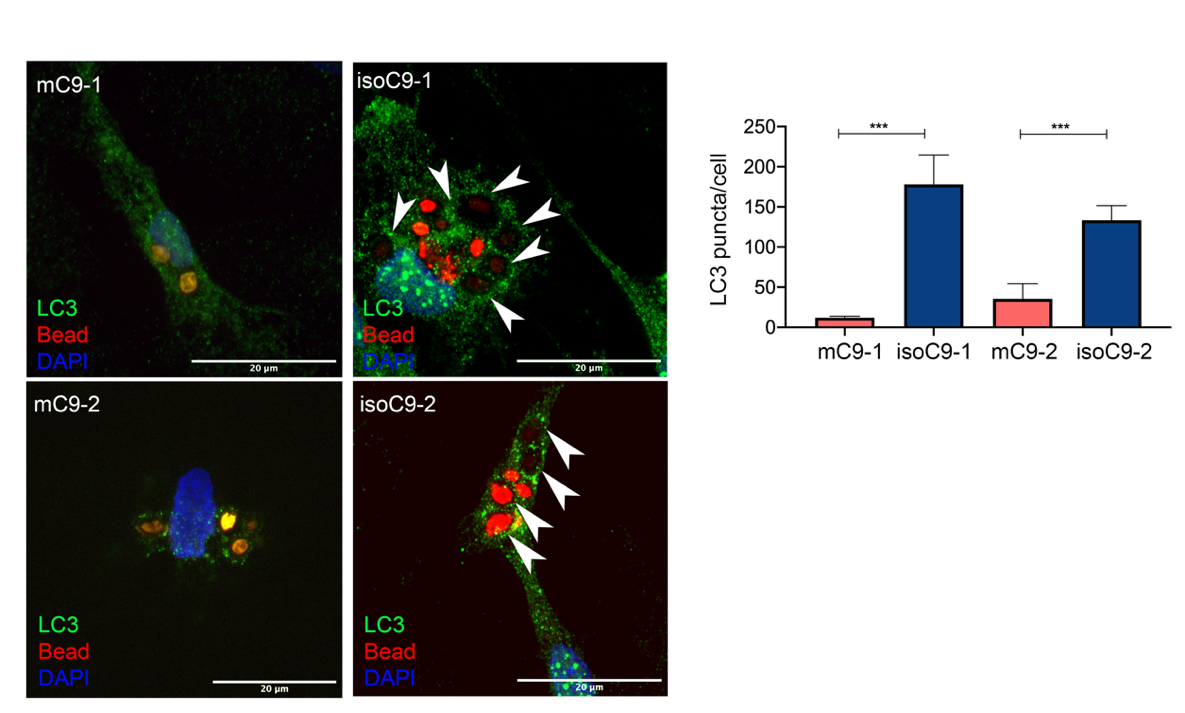

**Supplementary Fig 6- Appearance of LC3 puncta during zymosan bead uptake assay:** Zymosan uptake elicits the appearance of LC3 puncta in the isoC9-MG as opposed to mC9-MG, the number of LC3 puncta are quantified in graph (right), data represents mean+/-SD, statistical analysis was performed by two way ANOVA and Tukey’s multiple comparison test, N=3 (n=5) (*** p ≤ 0.001).

**Supplementary Fig :7 (Suppl. to Fig 4)**

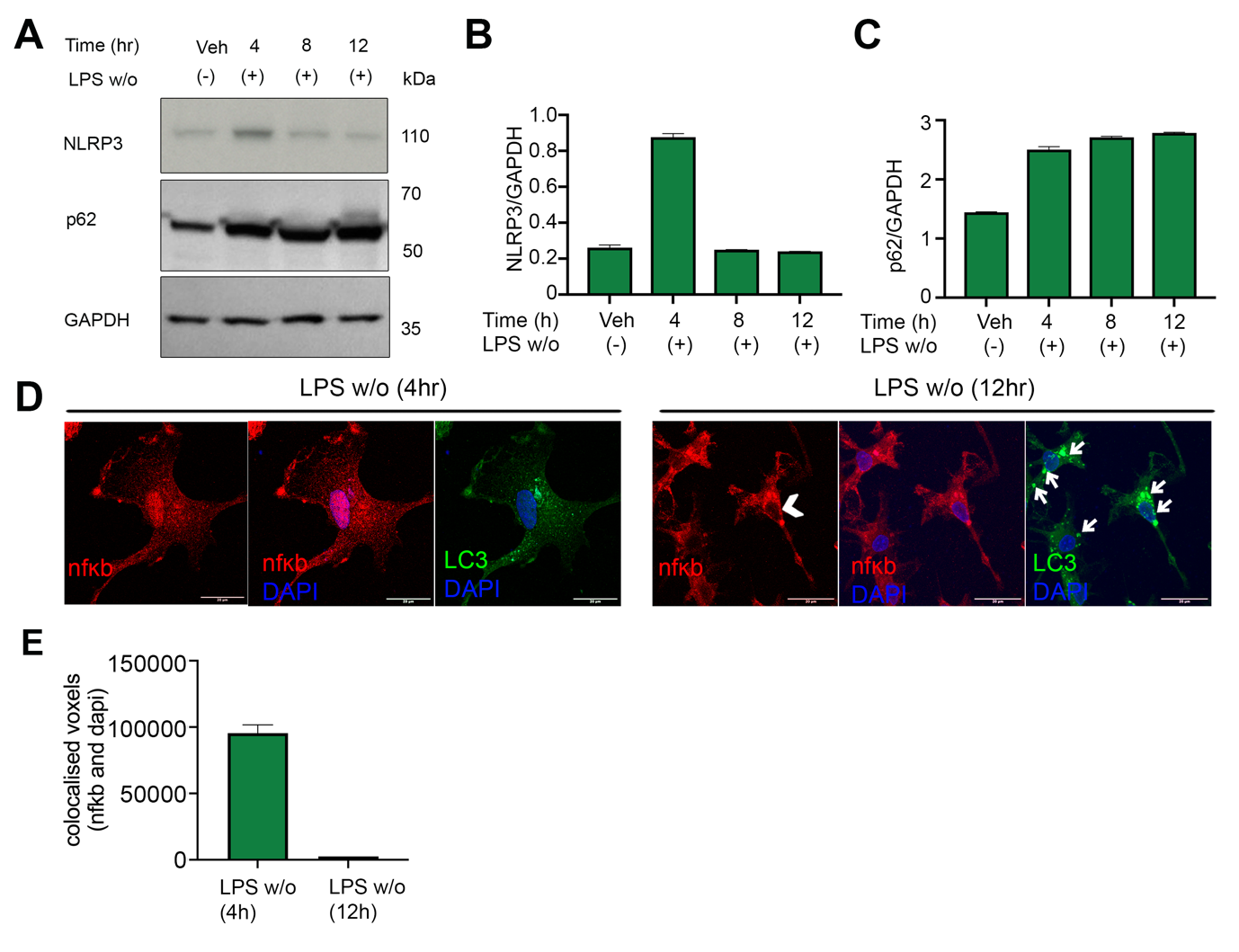

**Supplementary Fig 7 – Assessment of NF-κb signalling and NLRP3 activation following LPS stimulation in ctrl-MG (A)** Immunoblots showing differential immunoreactivity for NLRP3, p62 at 4h, 8h and 12h post LPS washout in ctrl-MG and vehicle treated cells (Veh) **(B, C)** Bar graphs showing densitometric values for NLRP3 and p62 normalised to GAPDH depicting a gradual decline of NLRP3 with a concomitant, reciprocal increase in p62 indicative of autophagy induction. **(D)** Representative images of immunofluorescence staining of NF-κB and LC3 demonstrating NF-κB signalling and autophagy activation at 4h and 12h post LPS removal. At 4h post LPS washout (left) ctrl-MG show nuclear localisation of NF-κB indicative of activated NF-κB signalling as opposed to the disappearance of NF-κB from the nucleus at 12 h post LPS washout indicative of an attenuation of NF-κB signalling, white arrow heads indicate the clearance of NF-κB from nucleus and white arrows indicate the appearance of LC3 puncta suggestive of autophagic activation at 12 h post LPS washout **(E)** demonstrates the quantification of nuclear localisation as denoted by the colocalised voxels of NF-κB and dapi; data is representative for 21 cells ; data represents mean +/-SD.

**Supplementary Fig 8 (Suppl. To Fig 4)**

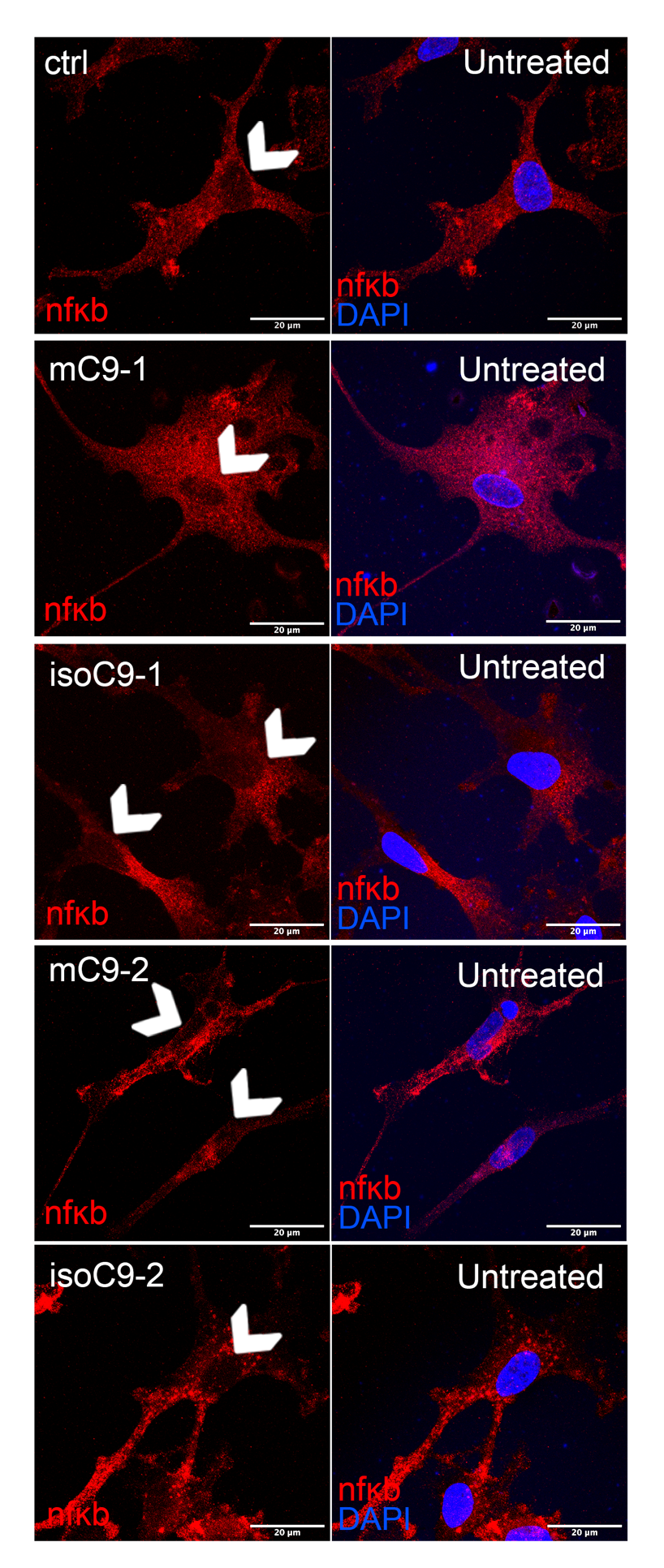

**Supplementary Fig 8:** **Localisation of NF-κB in ctrl-MG, mC9-MG and isoC9-MG in unstimulated condition:** Representative images of immunofluorescence staining showing the localisation of NF-κB at basal state across ctrl-MG, mC9 1-MG, isoC9 1-MG, mC9 2-MG and isoC9 2-MG. White arrow heads show the absence of nuclear nfkb. Scale bar=20um

**Supplementary Fig 9 (Suppl. to Fig 5)**

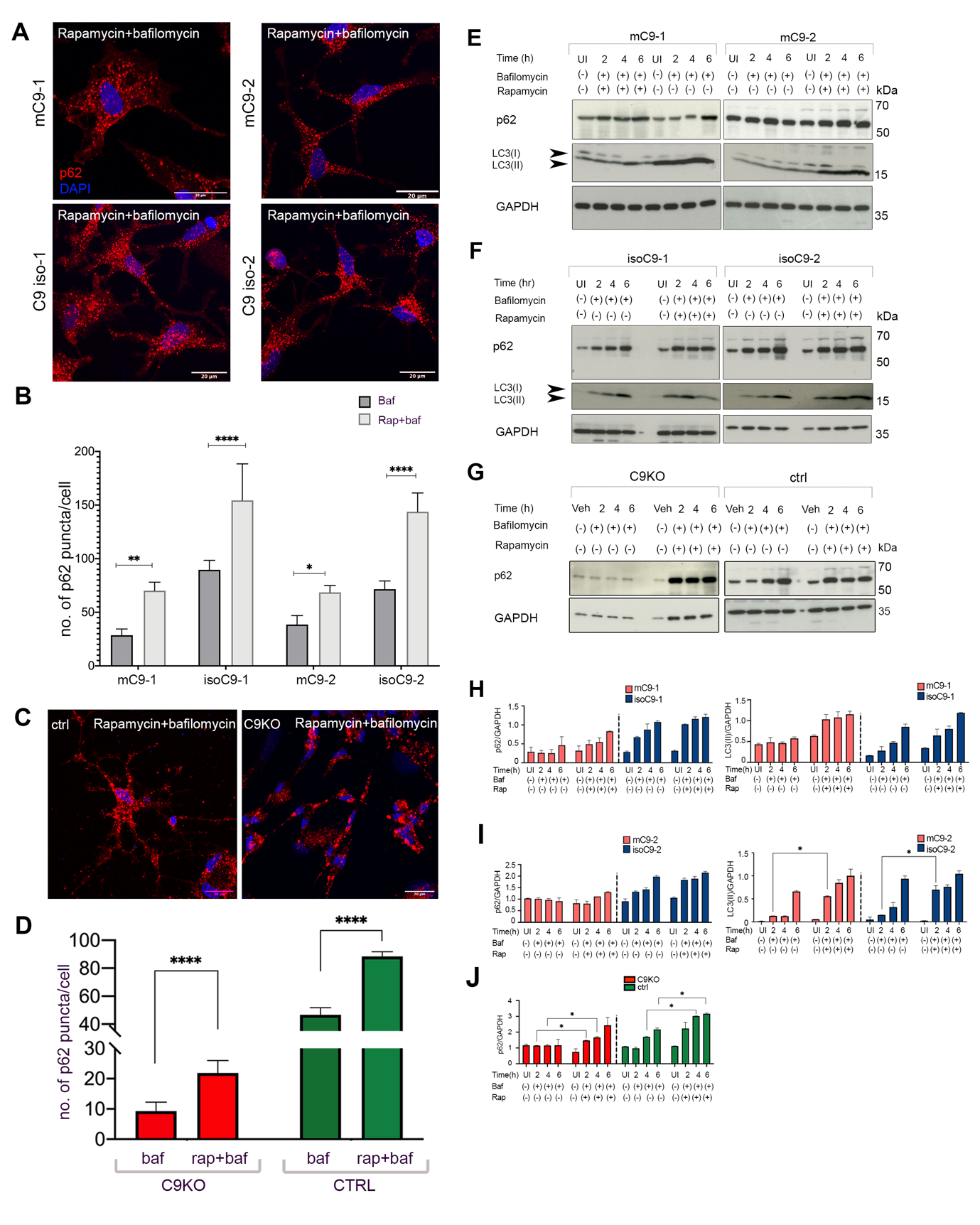

**Supplementary Fig 9: Rapamycin induces autophagy in mC9-MG, isoC9-MG, C9KO-MG and ctrl-MG**

**(A)** Representative images of immunofluorescence staining showing the increase in the number of p62(+) ve puncta in presence of 100nM rapamycin and 100nM bafilomycin for 6 hours in m*C9* 1-MG, m*C9* 2-MG and iso*C9* 1-MG and iso*C9*2-MG **(B)** Bar graph shows the quantification of p62 puncta from the immuno-fluorescence images across m*C9*-MG and iso*C9*-MG depicting an increase in the accumulation of puncta in the mC9 -MG upon treatment with rapamycin, Statistical analysis was performed using 2-way ANOVA and Tukey’s multiple comparison test, (* p ≤ 0.05; ** p ≤ 0.01 *** p ≤ 0.001;) **(C)** Representative images of immunofluorescence staining of C9KO-MG and ctrl-MG showing the increased number of p62+ve puncta post rapamycin treatment at 6h in C9KO-MG and ctrl-MG, these experiments were conducted in presence of 100nM bafilomycin. **(D)** Bar graph depicting the quantification of the number of p62 puncta in C9KO-MG and ctrl-MG post bafilomycin treatment in presence and absence of rapamycin, data is represented as mean +/-SD N=3 (n=10 cells were assessed across both genotypes). Statistical analysis was performed using two-way ANOVA and Tukey’s multiple comparison test, **** p ≤ 0.0001 **(E,F)** immuno-blots showing the increased turnover of p62 and LC3(II)in both pairs of mC9-MG and isoC9-MG when compared to rapamycin untreated condition and vehicle treated cells (Veh), indicative of enhanced autophagy induction **(G)** Immunoblots showing time dependent increase in the level of p62 in C9KO-MG and ctrl-MG following rapamycin treatment when compared to rapamycin untreated condition and vehicle treated cells (Veh), indicative of enhanced autophagy induction **(H,I)** Densitometric quantification of the level of p62, LC3 (II) over GAPDH shows the increment in the levels of p62 and LC3 post rapamycin treatment at 2h,4h,6h h in mC9 -MG, isoC9-MG, **(J)** Densitometric quantification of p62 levels depicting a time dependent increase of p62 level across C9KO-MG and ctrl-MG at 2h,4h,6h post rapamycin treatment.

**Supplementary Fig 10 (Suppl. To Fig 5)**

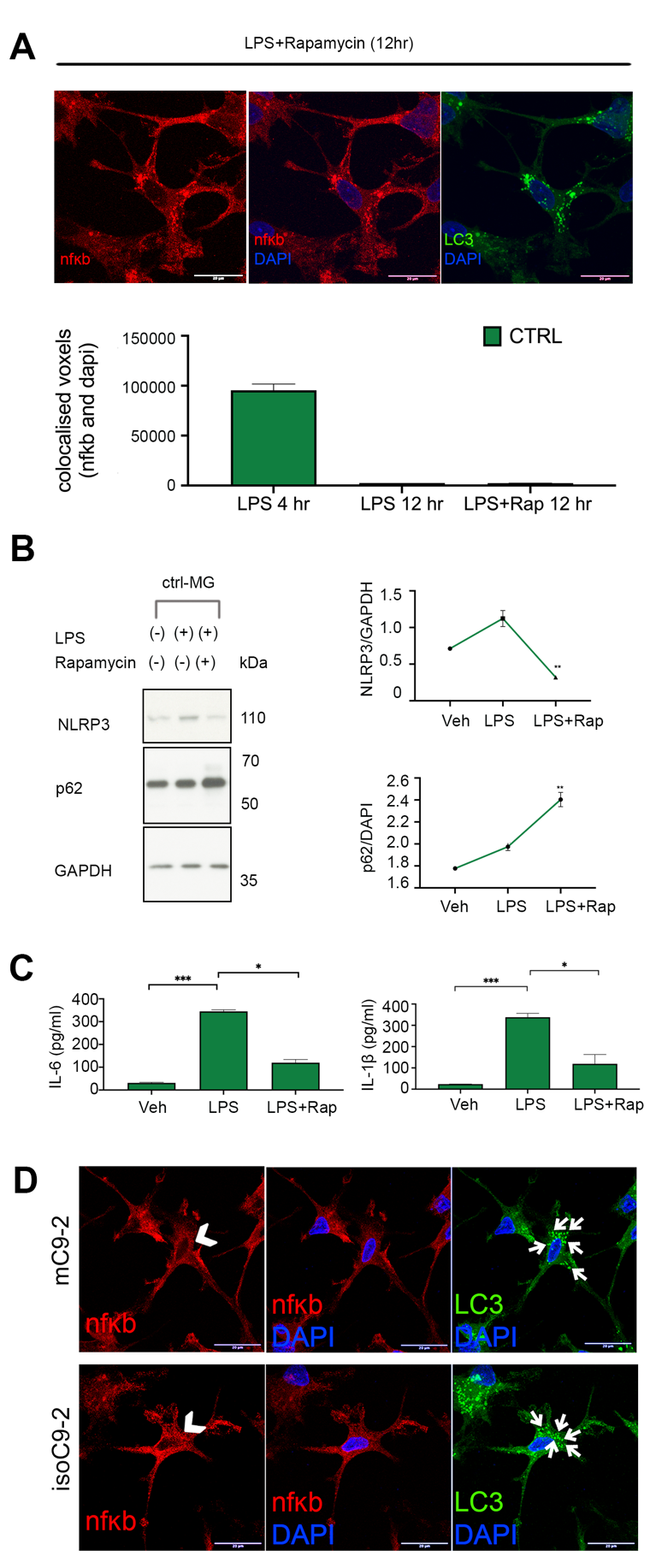

**Supplementary Fig 10: Rapamycin dampens the effect of LPS and enables the attenuation of nfkb signalling in ctrl-MG and mC9-MG (A)** Representative images of immunofluorescence staining showing the reduction of nuclear localisation of NF-κB in rapamycin-treated ctrl-MG, 12 hours post LPS washout, quantification of the colocalization of nfkb in the nucleus is shown in the bar graphs indicating a suppression of nfkb signalling as a result of rapamycin treatment. **(B)** Immunoblots and its quantification (right) showing an increase in p62 immunoreactivity and a reciprocal decline in NLRP3 immunoreactivity in presence of rapamycin at 12 h. p value was calculated using one-way ANOVA and multiple comparison analysis was performed using Tukey’s multiple comparison test, data represents mean+/-SD, N=3 (** p ≤ 0.01) **(C)** Cytokine ELISA of IL-6 and IL-1β demonstrates suppression of the production of pro-inflammatory cytokines in rapamycin treated ctrl-MG when compared to LPS only treated cells alongside vehicle treated cells (veh); p value was calculated using two-way ANOVA and multiple comparison analysis was performed using Tukey’s multiple comparison test, data represents mean+/-SD, N=3 (* p ≤ 0.05; *** p ≤ 0.001;). **(D)** immunofluorescence staining showing the reduction of nuclear localisation of NF-κB in rapamycin-treated m*C9*2-MG, demonstrating an attenuation of NF-κB signalling. The nuclear regions are indicated using white arrow heads and concurrent increase in appearance of LC3 puncta are highlighted using white arrows.

**Supplementary Fig: 11 (Suppl. Fig 6)**

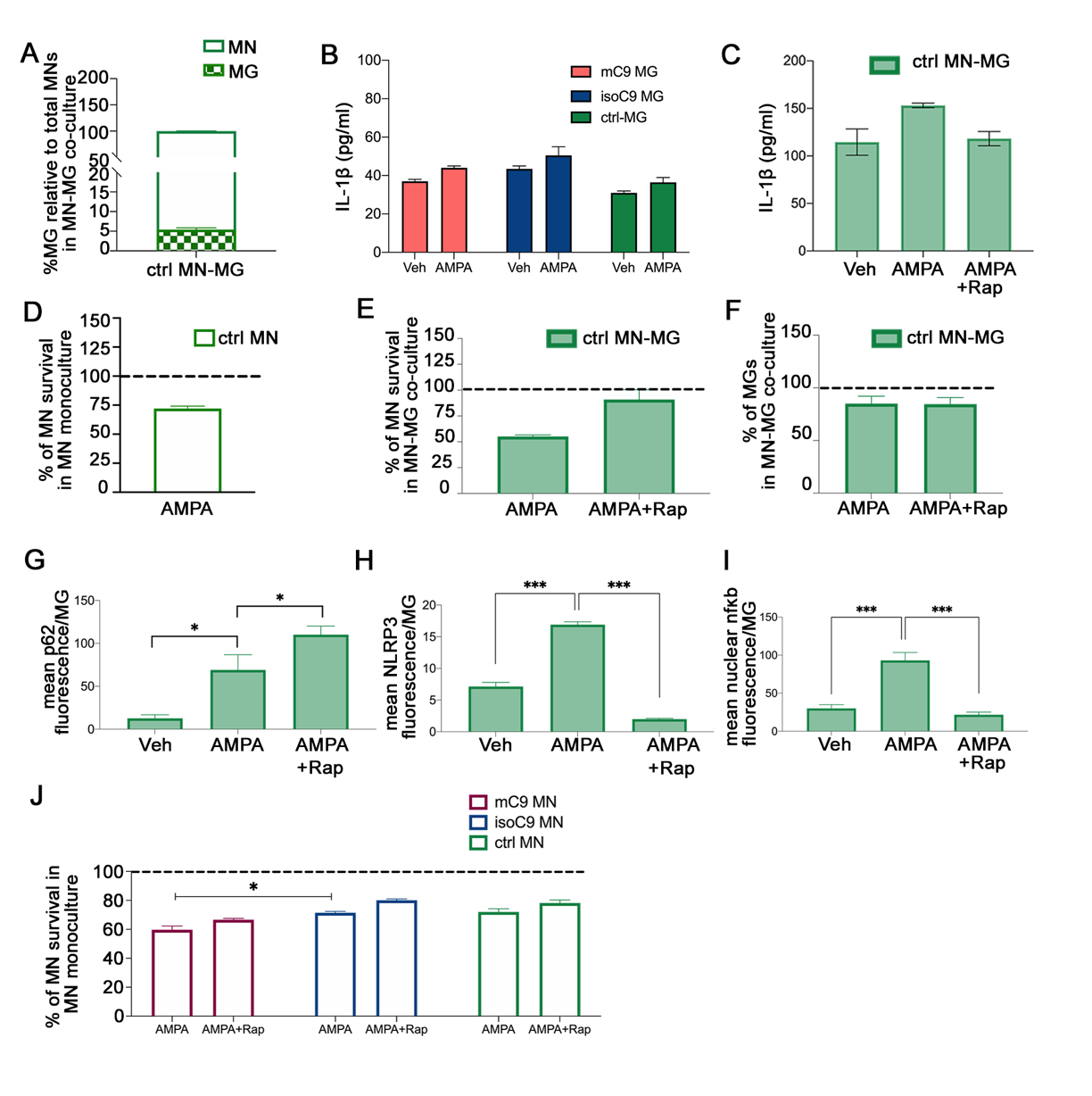

**Supplementary Fig 11: Assessment of neuronal death, microglial cytokine release, autophagy initiation, NF-κb signalling and NLRP3 activation following AMPA stimulation in ctrl MN-MG co-culture** (A) Graph representing the percentage of microglia (MG) relative to total number of motor neurons (MN) in ctrl MN-MG co-culture. Data are represented as mean +/- SD N=3 (B) Graph representing IL-1β release from monocultures of mC9-MG , isoC9-MG and ctrl-MG following AMPA challenge for 24 hours. Data are represented as mean +/- SD N=3 (C) Graph representing the production of IL1β in ctrl MN-MG co-cultures in vehicle treated condition and following AMPA challenge in absence and presence of rapamycin. Data are represented as mean +/- SD N=3 (D) Graph representing the percentage of motor neuron survival in ctrl-MN monoculture relative to their vehicle treated controls (represented by dashed line) following AMPA treatment for 24 hours. Data are represented as mean +/- SD N=3 (E) Graphs showing the percentage of motor neuron survival in ctrl MN-MG co-culture following AMPA treatment for 24 hours in absence and presence of rapamycin relative to their vehicle treated controls as represented by the dashed line (F) Graph representing the proportion of MGs in ctrl MN-MG co-culture relative to their vehicle treated condition (represented by the dashed line) following AMPA treatment for 24 hours in absence and presence of rapamycin. Data are represented as mean +/- SD N=3 (G,H,I) Graph representing the quantification of the mean fluorescence intensity of p62/NLRP3/nfκb per microglial cell in ctrl MN-MG co-culture across vehicle treated, AMPA treated conditions in absence and presence of rapamycin. 15 microglial cells from three biological replicates have been analysed per condition, data are represented as mean +/- SD N=3 (J) Graph showing percentage of motor neuron survival across mono-cultures of mC9-MN, isoC9-MN and ctrl-MN following AMPA treatment in absence and presence of rapamycin, vehicle treated condition is represented by the dashed line; data are represented as mean +/- SD N=3
